# Subcortically generated movements activate motor cortex during sleep and wake in rats through postnatal day 24

**DOI:** 10.1101/2025.01.26.634890

**Authors:** Madilyn R. Reid, Nicholas J. Sattler, James C. Dooley

**Affiliations:** Department of Biological Sciences, Purdue University, West Lafayette, Indiana; Purdue Institute for Integrative Neuroscience, Purdue University, West Lafayette, Indiana

**Author notes:** Corresponding author: James C. Dooley, Ph.D. These authors contributed equally (Madilyn R. Reid, Nicholas J. Sattler). Lead Contact: James C. Dooley.

**Keywords:** Primary motor cortex, development, sensorimotor, REM sleep, myoclonic twitching, red nucleus, rat

## Abstract

The development of motor control in primary motor cortex (M1) requires both movement and neural activity. In rats, cortical motor control first appears around postnatal day (P) 25, prior to which movements are generated by subcortical motor nuclei such as the red nucleus (RN). While these subcortical movements are thought to provide the activity that guides the development of M1, the types of movements that drive M1 activity and their somatotopic and temporal precision remain unknown. Here, we recorded activity in the forelimb region of M1 of P12–24 rats as they cycled between sleep and wake, and compared M1 and RN activity in P24 rats. At every age, M1 neurons showed somatopically precise activity to REM sleep twitches, along with strong responses to wake movements. From P12 to P24, the proportion of neurons that showed twitch-related activity decreased, twitch-related activity became more temporally refined, and a larger fraction of spikes occurred before movement onset. At P24, M1 showed less premovement activity than RN. Further, in contrast to the non-selective activity seen in M1, some RN neurons showed selective movement-related activity during wake, firing only during particular wake movements. These findings reveal that movement-related activity in M1 is somatotopically precise by P12 and temporally precise by P24. But M1 still lacks the strong premovement activity and selectivity characteristic of RN, suggesting that at P24, subcortical outputs remain the main drivers of M1’s movement-related activity.

**Significance Statement:** In adults, primary motor cortex (M1) is widely recognized for its role in initiating voluntary movements and guiding motor learning. However, during early development, our findings show that M1 processes activity related to self-generated limb movements but does not drive the production of those movements. Using simultaneous recordings from M1 and the red nucleus (RN) of developing rats, we show that although M1 shows movement-related activity during wake and sleep, RN remains the principal driver of behavior through at least postnatal day 24. These results refine our understanding of early movement-related activity and highlight the possibility that both sleep-related twitches and wake movements shape sensorimotor development by driving neural activity in M1 and RN.

## Introduction

In placental mammals, primary motor cortex (M1) is viewed as the key player in motor production and learning (see Peters et al., 2017). However, it is increasingly clear that motor cortex does not act alone, as coordinated movements necessitate activity from the entire brain, including the thalamus, basal ganglia, cerebellum, and brainstem (Kawai et al., 2015; Sauerbrei et al., 2020; Ruder et al., 2021; Inagaki et al., 2022; Lopez-Virgen et al., 2023; Yang et al., 2023). Yet in infancy, the contributions of these subcortical structures are impossible to ignore, as M1 is not yet capable of driving movement (Martin, 2005; Young et al., 2012; Williams and Martin, 2015; Singleton et al., 2021). Instead, in numerous mammals, including rats, anatomical and functional evidence shows that limb movements are generated by brainstem motor nuclei, with the midbrain’s red nucleus (RN) playing a critical role throughout this period (Ulfig and Chan, 2001; Rio-Bermudez et al., 2015; Williams and Martin, 2015).

However, M1’s activity is still important throughout infancy, as both M1 activity and ongoing movements are necessary for the development of cortical motor control (Chakrabarty and Martin, 2005; Martin et al., 2005), leading to the inference that early movements provide the activity that is necessary for M1’s development. M1 responds to sensory feedback from twitches—brief, stereotyped movements occurring during REM sleep—throughout early infancy (Tiriac et al., 2014; Tiriac and Blumberg, 2016). Starting around postnatal day (P) 12, M1 also responds to wake movements (Dooley and Blumberg, 2018; Glanz et al., 2021; Gómez et al., 2021). Intracortical microstimulation experiments indicate that M1’s motor maps first appear at P25 (Young et al., 2012; Singleton et al., 2021), providing M1 with nearly two weeks for ongoing movement-related activity to refine M1 circuits.

Thus, despite clear evidence that both twitches and wake movements pattern M1 activity early in life, it is not known how these responses evolve leading up to the development of motor control. This gap leaves unresolved questions about the somatotopic and temporal precision of the activity generated by twitches and wake movements. Moreover, the extent—or indeed even the existence—of premovement activity in M1 from P12 to P25 remains unknown, as prior studies were conducted under anesthesia, and thus cannot capture how such activity unfolds in naturally behaving animals. Finally, how movement-related activity in M1 compares to that of RN—the subcortical motor center presumed to be responsible for limb movements during this transition—has not been examined. Addressing these questions is essential for understanding how subcortical circuits sculpt M1 during early development, enabling its eventual role in motor control.

Here, we test these questions by performing extracellular recordings in the forelimb representation of M1 in unanesthetized rats spanning P12 to P24 as they cycle between non-REM (NREM) sleep, REM sleep, and wake. By analyzing movement-related activity across sleep and wake, we show that M1 neurons respond to REM-sleep associated twitches and wake movements at all ages (P12 to P24). Taking advantage of the discreteness of twitches, we show that twitches of body parts other than the forelimbs fail to drive activity in the forelimb region of M1, illustrating that movement-related activity in M1 is somatotopically refined by P12. Whereas M1 exhibited more premovement spiking by P24, it remained unclear whether this activity reflects emerging motor output—a question we addressed by comparing M1 directly to RN in P24 rats. We found that activity in RN precedes twitches and wake movements to a significantly greater extent than M1. Further, we found a subset of neurons in RN fire during specific limb movements, whereas neurons in M1 were more indiscriminate in their movement-related activity. Altogether, our results suggest that premovement activity is emerging in M1 by P24 but remains less pronounced than in RN. Together with the selective responses of RN neuron’s to different wake movements, these data highlight the continued dominance of subcortical circuits for motor control at this stage.

## Materials and Methods

### Code Accessibility

All original code has been deposited at https://github.com/jcdooley/XXX and is publicly available as of the date of publication. Any additional information required to reanalyze the data reported in this paper is available from the corresponding author upon request.

### Experimental model and subject details

For the first experiment, neurophysiological recordings of M1 were obtained from Sprague-Dawley Norway rats (*Rattus norvegicus*) at P12 (N = 9 pups; 30.8 ± 2.4 g; 3 male), P15-16 (hereafter P16; N = 12 pups; 43.7 ± 3.0 g; 8 male), P18-P20 (hereafter P20; N = 12 pups; 56.1 ± 4.4 g; 7 male), and P22-P24 (hereafter P24; N = 8 pups; 68.5 ± 4.7 g; 6 male). Animals within the same age group were always from different litters. Although the focus here is on M1 activity, recordings were performed in the forelimb region of M1 and either the ventral lateral or the ventral posterior nucleus of thalamus. The thalamic data has been published and described previously (Dooley et al., 2021). For the second experiment, dual neurophysiological recordings were performed in M1 and the RN at P24 (N = 7 pups, 65.8 ± 5.2 g; 3 male). To be included in the analysis, animals needed more than 40 right forelimb twitches and at least one neuron had to be responsive to right forelimb twitches. This criterion led to the exclusion of 2 animals.

Pups were born and raised in standard laboratory cages (48 × 20 × 26 cm) in a temperature- and humidity-controlled room on a 12:12 light-dark cycle, with food and water available ad libitum. The day of birth was considered P0 and litters were culled to eight pups by P3. All pups had at least four littermates until P12 and at least two littermates on the day that neurophysiological recordings were performed. No pups were weaned before testing. All experiments were conducted in accordance with the National Institutes of Health (NIH) Guide for the Care and Use of Laboratory Animals (NIH Publication No. 80–23) and were approved by the Institutional Animal Care and Use Committees at the University of Iowa and Purdue University.

### Method Details

*Surgery.* As described previously (Dooley et al., 2021), a pup with a healthy body weight and (for P12 pups) a visible milk band was removed from the litter and anesthetized with isoflurane gas (3–5%; Phoenix Pharmaceuticals, Burlingame, CA). The hair on the top of the head was shaved, and care was taken to ensure that the vibrissae were intact. For sleep/wake state determination, two custom-made bipolar hook electrodes (0.002-inch diameter, epoxy coated; California Fine Wire, Grover Beach, CA) were inserted into the nuchal and biceps muscles contralateral to the neural recordings. Carprofen (5 mg/kg SC; Putney, Portland, ME) was administered as an anti-inflammatory analgesic.

The skin above the skull was carefully removed and an analgesic (bupivicane; Pfizer, New York, NY) was applied topically to the skull. The skull was then dried with bleach. For P20 and P24 rats, any bleeding around the skull was cauterized to ensure that it was completely dry prior to the application of a headplate. Vetbond (3M, Minneapolis, MN) was then applied to the skin surrounding the incision, and a custom-manufactured head-plate (Neurotar, Helsinki, Finland) was secured to the skull using cyanoacrylate adhesive.

A trephine drill (1.8 mm; Fine Science Tools, Foster City, CA) was used to drill a hole into the skull above the forelimb representation of M1 (0.5 mm anterior and 2.2–2.5 mm lateral to bregma). When thalamic recordings were also performed (as reported in Dooley et al., 2021), a second hole was drilled above thalamus (2.0–2.8 mm caudal to bregma, 2.2– 2.5 mm lateral to the sagittal suture). In Experiment 2, when RN recordings were performed, a second hole was drilled above RN (5.2 mm caudal to bregma, 0.4 to 0.8 mm lateral to the sagittal suture). A small amount of peanut oil was applied to the dura to prevent desiccation of the underlying tissue. This surgical procedure lasted approximately 30 min.

While recovering from anesthesia, the pup was secured to a custom-made head-fixation plate secured to a Mobile HomeCage (NTR000289-01; Neurotar). The height of the pup’s head off the floor was adjusted depending on its age so that, when the pup was exhibiting atonia during REM sleep, its body rested on its elbows (P12: 35 mm; P16: 38 mm; P20 and P24: 40 mm). Electromyography (EMG) wires were carefully secured behind the pup’s back to ensure that they would not get tangled. Pups recovered from anesthesia within 15 minutes and were acclimated to the head-fix apparatus for 1 to 2.5 hours. This period of acclimation allowed the recording electrodes to stabilize and the return of normal spiking activity (Domínguez et al., 2021). When recording began, all pups were exhibiting typical behavioral and electrophysiological features of sleep and wake (i.e., twitching, grooming, locomotion).

*Recording Environment.* Electrophysiological recordings were performed in a Faraday cage illuminated by flicker-free red LEDs (630 nm; Waveform Lighting, Vancouver, WA). Continuous white noise (70 dB) was present throughout the recording session. To maximize REM sleep, the room’s temperature was maintained between 26.5 and 29° C (Szymusiak and Satinoff, 1981). The animal’s head was positioned away from the entry to the room so that it could not see the experimenter entering or leaving, which was infrequent. For the comfort of both the experimenter and the subject, the experimenter was outside the room and monitoring the animal and data acquisition remotely.

*Electrophysiological Recordings.* The nuchal and biceps EMG electrodes were connected to the analogue inputs of a Lab Rat LR-10 acquisition system (Tucker Davis Technologies, Gainesville, FL). The EMG signals were sampled at approximately 1.5 kHz and high-pass filtered at 300 Hz.

Before insertion, we coated a 16-channel silicon depth electrode (A1×16-3mm-100-177-A16) with fluorescent DiI (Vybrant DiI Cell-Labeling Solution; Life Technologies, Grand Island, NY). Electrodes were inserted using a multiprobe manipulator (New Scale Technologies; Victor, NY) controlled by an Xbox controller (Microsoft, Redmond, WA). The electrode was inserted into cortex until all electrode sites were beneath the dura (approximately 1500 μm). A chlorinated Ag/Ag-Cl wire (0.25 mm diameter; Medwire, Mt. Vernon, NY) inserted into occipital cortex contralateral to the cortical recording sites served as both reference and ground. Neural signals were sampled at approximately 25 kHz. A high-pass filter (0.1 Hz) and a harmonic notch filter (60, 120, and 180 Hz) were applied.

Electrophysiological data were acquired continuously for 3-6 h using SynapseLite (Tucker Davis Technologies). The Mobile HomeCage enabled stable head-fixation throughout the recording session while rats could freely locomote, groom, and sleep.

*Video Collection and Synchronization.* As described previously (Dooley et al., 2020), video data were synchronized to electrophysiological recordings so that we could quantify movements and behavioral state. Rats were surrounded by a clear enclosure within the Mobile HomeCage, enabling unimpeded visual access. The video was recorded using a single Blackfly-S camera (FLIR Integrated Systems; Wilsonville, Oregon) positioned at a 45° angle (relative to the pup’s head) and centered on the right forelimb, contralateral to the M1 recordings. This angle provided an unobstructed view of the right whiskers, forelimb, hindlimb, and tail. Video was collected in SpinView (FLIR Integrated Systems) at 100 frames/s, with a 7 ms exposure time and 720-x 540-pixel resolution.

Video frames were synchronized to the electrophysiological record using an external time-locking stimulus. A red LED, controlled by SynapseLite (Tucker Davis Technologies), was in view of the camera and pulsed every 3 s for a duration of 100 ms. Custom MATLAB scripts determined the number of frames between each LED pulse to check for dropped frames. Although infrequent, when the number of frames between pulses was not equal to 300 (3 s inter-pulse interval x 100 frames/s), a “dummy frame” was inserted in that location. This ensured that the video and electrophysiological data were synchronized to within one frame (10 ms) throughout data collection.

*Histology.* At the end of the recording session, the pup was euthanized with ketamine/xylazine (10:1; >0.08 mg/kg) and perfused with 0.1 M phosphate-buffered saline (PBS) followed by 4% paraformaldehyde (PFA). The brain was extracted and post-fixed in 4% PFA for at least 24 h and was then transferred to a 30% sucrose solution at least 24 h before sectioning.

To confirm the electrode’s location in M1, the neocortex was either flattened and sectioned tangentially to the pial surface or sectioned coronally. When sectioned tangentially, the right cortical hemisphere was dissected from the subcortical tissue and flattened between two glass slides (separated using a 1.5 mm spacer) for 5-15 min. Small weights (10 g) applied light pressure to the upper slide. Regardless of the plane of section, the tissue was sectioned at 80 µm. Wet-mounted sections were imaged at 2.5x using a fluorescent microscope and digital camera (Leica Microsystems, Buffalo Grove, IL) to identify the location of the DiI.

Cortical sections were stained for cytochrome oxidase (CO), which reliably delineates primary sensory areas in cortex at these ages (Seelke et al., 2012). Briefly, cytochrome C (3 mg per 10 mL solution; Sigma-Aldrich), catalase (2 mg per 10 mL solution; Sigma-Aldrich) and 3,3’-diaminobenzidine tetrahydrochloride (DAB; 5 mg per 10 mL solution; Spectrum, Henderson, NV) were dissolved in a 1:1 dilution of PB-H2O and distilled water. Sections were developed in well plates on a shaker at 35–40°C at approximately 100 rotations per min for 3–6 h, after which they were washed in PBS, mounted onto glass slides, and allowed at least 48 h to dry. Once dry, sections were placed in citrus clearing solvent (Richard-Allan Scientific) and cover slipped.

RN sections were stained using cresyl violet, which reliably delineates the large cell bodies in RN from the surrounding tissue (Rio-Bermudez et al., 2015). On mounted gelatin-coated slides, sections were dehydrated via submersion in increasing concentrations of ethanol in distilled water for 5-min each. Sections were then defatted using two 5-min washes of citrus clearing solution (Richard-Allan Scientific), followed by a rinse in 100% ethanol. Sections were then stained in a 0.1% cresyl violet solution for 5-10 min, rinsed in distilled water, and immersed in a differentiation solution (70% ethanol with 0.1-0.5% acetic acid) until cell nuclei were maximally differentiated. Finally, sections were fully dehydrated in 100% ethanol for 5-min, followed by 5-min in 2 subsequent rinses of citrus clearing solvent, and cover slipped.

Stained sections were imaged at 2.5x or 5x magnification. Multiple images were combined into a single composite image (Microsoft Image Composite Editor; Microsoft, Redmond, WA) and the electrode location were visualized in relation to areal, nuclear, and laminar boundaries of the stained tissue. M1 recordings were included in subsequent analyses when they were between 0 and 1.2 mm medial to the forelimb representation of S1. RN recordings were included in subsequent analyses when the electrode track went through the large cell bodies of RN.

*Local Field Potential.* For all local field potential (LFP) analyses, the raw neural signal was smoothed using a moving Gaussian kernel with a half-width of 0.5 ms and then down sampled to ∼1000 Hz.

*Classification of Behavioral State.* As described previously, behavior, nuchal EMG signals, and cortical LFPs were used to identify periods of wake, REM sleep, and NREM sleep (Dooley et al., 2021). Wake was characterized by periods of high nuchal muscle tone and wake-related behaviors (e.g., locomotion, grooming). REM sleep was characterized by the occurrence of myoclonic twitches against a background of muscle atonia. At P16 and later, REM sleep was also accompanied by the presence of continuous theta oscillations in thalamic and RN recordings. During NREM sleep, we observed high cortical delta power, behavioral quiescence, and moderate nuchal muscle tone, although periods of high delta power were sometimes accompanied by nuchal muscle atonia. Periods not assigned as REM or NREM sleep were designated as wake. Active wake was defined as that part of the wake period that was less than 3 s following an ROI-defined period of movement (see below).

*ROI Movement Classification.* As described previously (Dooley et al., 2021), to quantify periods of movement, we used custom MATLAB scripts to detect frame-by-frame changes in pixel intensity within regions-of-interest (ROIs). For each ROI, the number of pixels exhibiting an intensity change > 5% was summed for each frame, producing a 100 Hz time series of movement-related pixel changes. This automated ROI-based detection was used to identify candidate movements (twitches or wake movements), which were then visually inspected with synchronized video to confirm the presence of a twitch or wake movement and to verify the involved body part.

For twitches, the movement timeseries for a ROI containing the body part of interest (e.g., forelimb) was visualized alongside the video using Spike2 (Version 8; Cambridge

Electronic Design, Cambridge, UK). Peaks in the time series were confirmed to be twitches of the appropriate body part. Twitch onset was defined as the first frame in which movement occurred (using the ROI timeseries data described above). For the forelimb, hindlimb, and tail, this method effectively counted every visible twitch, since even twitches in rapid succession have distinct onset and offset times. However, for whisker twitches, alternating protractions and retractions did not always have clear temporal boundaries, thus making it difficult to determine when one whisker twitch ended, and another began. In these instances, only the first whisker twitch in a series of twitches was counted.

For wake movements, an ROI of the entire animal was used because forelimb position during wake was often too variable for a fixed, body-part-specific ROI. Candidate wake movements were only included in subsequent analyses when the movement involved the forelimb, and the movement’s initiation time was defined as the first frame in which the forelimb moved independently. To ensure that each labeled wake movement represented a clear transition from quiescence to movement, only the first movement in a bout was marked, with subsequent movements labeled only if separated by at least 500 ms of behavioral quiescence.

### Quantification and statistical Analysis

*Spike Sorting.* SynapseLite files were converted to binary files using custom MATLAB scripts and sorted with Kilosort 2.0 (Pachitariu et al., 2016). Briefly, data were whitened (covariance-standardized) and band-pass filtered (300-5000 Hz) before spike detection. Template-matching was implemented to sort the event waveforms into clusters. The first-pass spike detection threshold was set to six standard deviations below the mean and the second-pass threshold was set to five standard deviations below the mean. The minimum allowable firing rate was set to 0.01 spikes/s and the bin size for template detection was set to 262,400 sample points, or approximately 11 s. All other Kilosort parameters were left at their default values.

Clusters were visualized and sorted in Phy2 (Rossant and Harris, 2013). Putative single units had spike waveforms that reliably fit within a well-isolated waveform template, appeared in a principal component analysis as a distinct cluster, and had an auto-correlogram with a decreased firing rate at a time lag of 0 (indicative of a unit’s refractory period).

Clusters meeting the first two criteria but not the third were considered multi-units and were discarded from analysis. Any putative unit with a waveform template indicative of electrical noise, a firing rate < 0.01 spikes/s, or amplitude drift across the recording period was discarded. Cross-correlograms of all single- and multi-units on nearby channels were compared. If cross-correlograms and auto-correlograms were similar, clusters were merged and (if appropriate) reclassified as multi-units.

*Determination of Twitch- and Wake-Movement-Responsiveness.* All analyses of neural data were performed in MATLAB using custom-written scripts. The relation between neural activity and twitches or wake movements was assessed as follows. First, all twitches of each body part (forelimb, hindlimb, whiskers, tail) were behaviorally scored. For each body part that had more than 20 twitches, perievent histograms of neural activity were constructed (window size: −3 to 3 s; bin size: 10 ms). Next, we determined the mean baseline firing rate from −3 to −0.5 s. Finally, we z-scored the perievent histograms by subtracting the baseline from the raw perievent histograms and dividing this value by the standard deviation of the baseline. A neuron was considered “responsive” to twitches if it showed a clear peak with a z-score of at least 3.5 for at least one body part. A neuron was considered “responsive” to wake movements using the same criteria, but with perievent histograms triggered on wake movements. All other neurons were classified as “non-responsive.”

To accurately determine a neuron’s preferred somatotopic response, we only analyzed twitches for a given body part that were separated from twitches of other body parts by at least 100 ms. The preferred body part for each neuron was the body part with the largest peak in the z-scored perievent histogram. At all ages, neurons in the forelimb region of M1 responded to right forelimb twitches, not right hindlimb, tail, or whisker twitches.

*ROI-based Movement Analysis.* For each movement type (twitches of different body parts and wake movements), we calculated the average displacement produced by that movement for each pup. To ensure subsequent twitches (of the same or different body parts) did not influence the average displacement of a twitch, for this analysis, we used the same subset of twitch triggers used in the somatotopy analysis above (i.e., twitches with inter-twitch-intervals of at least 100 ms). Median average displacement was calculated over all triggered twitches (or wake movements) at each timepoint. Because the size of the ROI used to assess movements varied across age and body part, median values for each animal were always normalized and then averaged.

*Movement kinematics.* For the M1-RN dataset, a point tracking the centroid position of the palm of the left forelimb and radiocarpal joint of the wrist for the right forelimb was used to quantify movement trajectories using DeepLabCut (Mathis et al., 2018). Manually labeled frames taken across periods of REM sleep and wake adjacent to movement from all animals were used to initially train the network. Frames with poor marker estimates from newly analyzed videos were then additionally labeled and used to re-train the network until mean likelihoods of 0.85 were achieved across previously detected twitches and wake movements. For all videos, 1 mm = 5.75 pixels.

*Movement angle and amplitude.* Using the previously established movement initiation times, the movement initiation location was defined as the position of the limb on the frame of movement onset. Using this point as the origin, we calculated the amplitude and angle of the limb at every timepoint for 150 ms for twitches, and 500 ms for wake movements. Movement amplitude was then defined as the limb’s peak displacement within the window, and the movement angle was defined as the limb’s angle at peak displacement.

*Width at Half-Height.* To measure width at half-height, the neural data were first smoothed using a 5-bin kernel and then upscaled from 10 ms bins to 1 ms bins using the interp1() function in MATLAB. The data were then normalized so that the baseline was equal to 0 and the peak was set to 1. The width at half-height (in ms) was calculated as the number of continuous bins greater than 0.5 around the peak of the normalized data.

*Proportion of spikes before movement onset.* For twitches, the proportion of spikes before movement onset was calculated by approximating the integral of z-scored perievent histograms triggered on twitch onset using the trapz() function in MATLAB. First, a window was selected that was inclusive of the period of elevated activity for the neurons being analyzed (−0.2 to 0.75 s for the M1 development analysis, and −0.2 to 0.2 s for the M1-RN analysis). The proportion of activity before twitch onset was calculated by dividing the integral from −0.2 to 0 s by the integral of the entire window.

For wake movements, the same method was used, with two adjustments. First, because the baseline period for wake movements was contaminated by prior movement, an adjusted baseline was used, defined as the minimum in the preceding 500 ms of a perievent histogram smoothed using a 20-bin kernel. Second, the window began at that minimum for each neuron and extended for 1 s after wake movement onset.

For three neurons from the twitch analyses and two neurons for the wake analysis, a negative value in the numerator produced a nonsensical negative proportion of spikes preceding twitch onset. Thus, for these neurons the proportion was set to 0.

*Coefficient of variation of neural response.* The coefficient of variation was calculated from the number of spikes within a window that included twitch- and wake-movement-related neural activity (−70 to 70 ms for twitches; −100 to 500 ms for wake movements). For each neuron, the standard deviation of the spikes was divided by the mean number of spikes within that window.

*Statistical Analyses.* All statistical tests were performed using MATLAB. Alpha was set at 0.05 for all analyses; when appropriate, the Bonferroni procedure was used to correct for multiple comparisons. Unless otherwise stated, mean data are always reported with their standard error of the mean (SEM). Data were tested for significance using a one-way ANOVA, two-way ANOVA, or t test. Categorical data were tested for significance using a Chi-squared test. Violin plots were constructed absent outliers, which were determined using the isoutlier() function in MATLAB. This function excludes data points that are more than three scaled median-absolute-deviations from the median.

## Results

We asked whether movement-related activity in M1 across sleep and wake shows developmental changes from P12 to P24, a period just before the onset of cortical motor control. Neural recordings were performed in the Mobile HomeCage, which permits locomotion under head fixation (**Figure 1A**) in a warm recording environment (∼29 °C) that promotes REM sleep (Szymusiak and Satinoff, 1981). Together, these features allowed us to record significant periods of REM and NREM sleep on the day of surgery, facilitating both REM sleep twitches and wake movements. Extracellular activity was recorded in the forelimb region of M1 (P12: N = 9 pups, 121 neurons; P16: N = 12 pups, 177 neurons; P20: N = 12 pups, 172 neurons; P24: N = 8 pups, 87 neurons; **Figure 1C, D**). Neural activity, electromyographic activity of the nuchal and biceps muscles, and high-speed video (100 frames/s) were recorded continuously for 3-6 h, and representative data from each behavioral state and each age can be seen in **Figure S1**. All rats cycled between sleep and wake throughout the recording sessions. Recordings were performed entirely during the lights-on period (1000 to 1800 hours). As expected, the percentage of time spent in REM sleep decreased across age, whereas the percentage of time spent in quiet wake increased (**Figure 1B**; Jouvet-Mounier et al., 1969).

**Figure 1.**
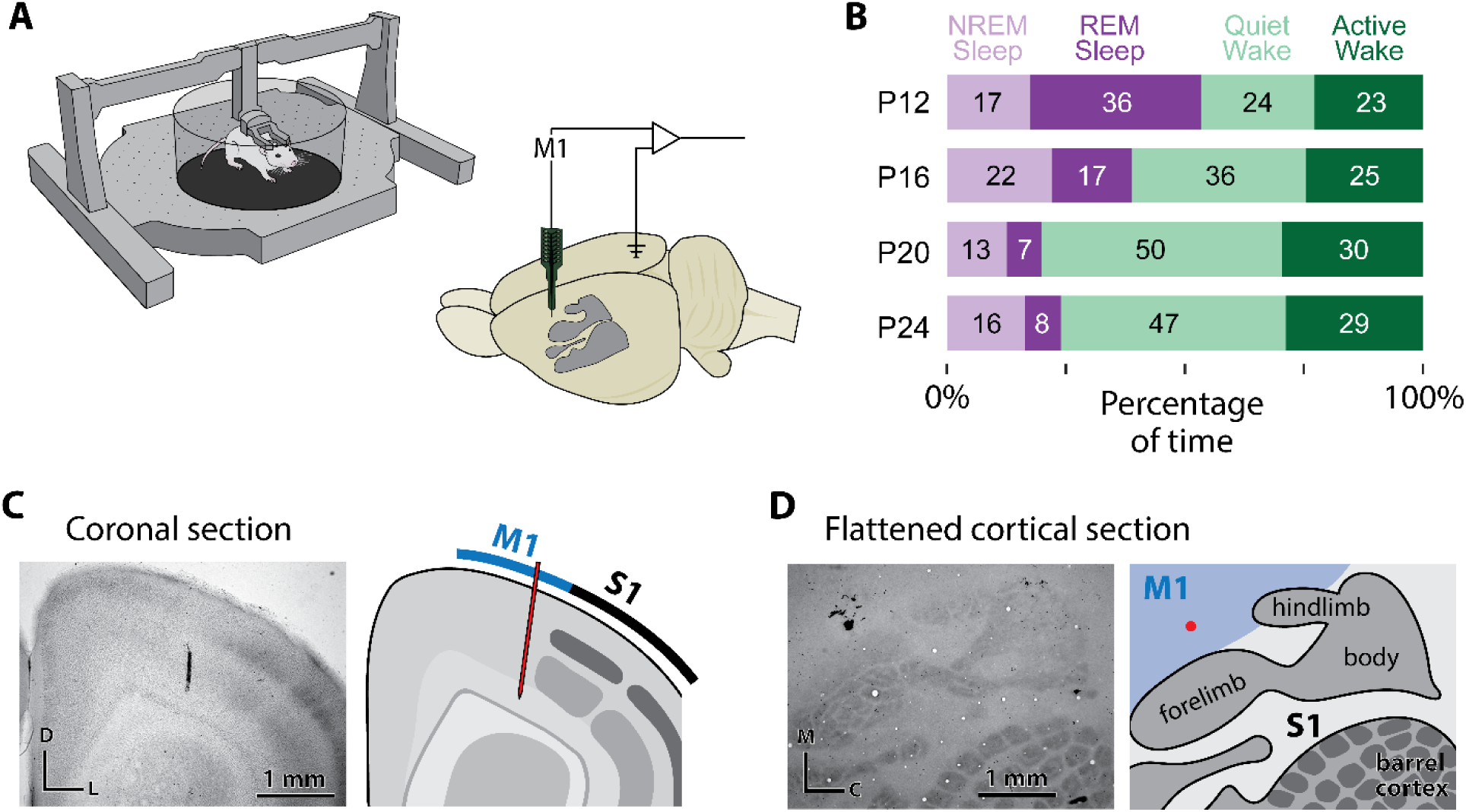
Recording environment, behavioral state quantification, and M1 recording verification. **(A)** Left: An illustration of the Mobile HomeCage used to record behavioral and electrophysiological data from head-fixed rat pups across sleep and wake. Right: A schematic of the rat brain with the primary somatosensory cortex (S1) shaded in gray, highlighting the approximate recording location in M1 and the reference/ground in the contralateral occipital cortex. **(B)** Mean percentage of time spent in NREM sleep (light purple), REM sleep (dark purple), quiet wake (light green), and active wake (dark green) at each age. Numbers in each bar indicate the mean percentage of total recording time for that state across all animals. **(C)** Representative coronal section stained for cytochrome oxidase showing the DiI-labeled electrode track in M1. Borders of primary somatosensory cortex are shown in gray; the DiI-labeled track is visible in the forelimb region of M1. **(D)** Representative flattened cortical section stained for cytochrome oxidase showing the DiI-labeled electrode track in M1. Conventions as in **C**.

We first asked whether overall M1 firing rates change across development or differ between behavioral states. M1 activity varied significantly by age during wake (F_(3, 553)_ = 6.51; p < 0.0005), NREM sleep (F_(3, 433)_ = 8.2; p < 0.0001), and REM sleep (F_(3, 553)_ = 3.56; p < 0.05; **Figure S2A-C**). However, there was not a general trend towards increased or decreased activity for REM sleep or wake, although NREM sleep did generally show an increase in activity form P12 to P24.

Next, we focused on directly comparing baseline neural activity during wake and REM sleep, the two behavioral states where movements are produced. We observed more activity during REM sleep than wake at P12 (t_120_ = 9.278, p < 0.0001), P16 (t_176_ = 6.55, p < 0.0001), and P20 (t_171_ = 8.23, p < 0.0001), but not at P24 (t_86_ = 1.20, p = 0.23; **Figure S2D**). The lack of a REM–wake difference at P24 parallels findings in adults, where activity in M1 is typically comparable, or even reduced, during REM sleep (Aime et al., 2022; Brécier et al., 2022). By contrast, the increased activity during REM sleep compared to wake observed at P12–P20 aligns with prior reports in younger pups (Tiriac and Blumberg, 2016; Dooley and Blumberg, 2018). Together, these results suggest that the infantile state, with REM sleep showing more neural activity than wake, appears to disappear between P20 and P24, a finding that, to our knowledge, has not been previously reported.

### M1 shows somatotopically precise movement-related activity from P12 to P24

Over half of M1 neurons responded to forelimb movements during wake and/or forelimb twitches during REM sleep (P12: 77%; P16: 74%; P20: 55%, P24: 65%; **Figure 2A**). At P12, a greater proportion of M1 neurons responded to twitches than wake movements. However, at P16 and beyond, more M1 neurons responded to wake movements than twitches, and only a small minority of neurons that were twitch responsive were not wake responsive (**Figure 2B**). This is not surprising, as wake movements are much longer in duration than twitches, and unlike twitches, involve the simultaneous activation of numerous muscles. Still, at all ages examined, over a quarter of movement-responsive neurons in M1 responded to twitches. Representative twitch responsive neurons at each age can be seen in **Figure 2C**.

**Figure 2.**
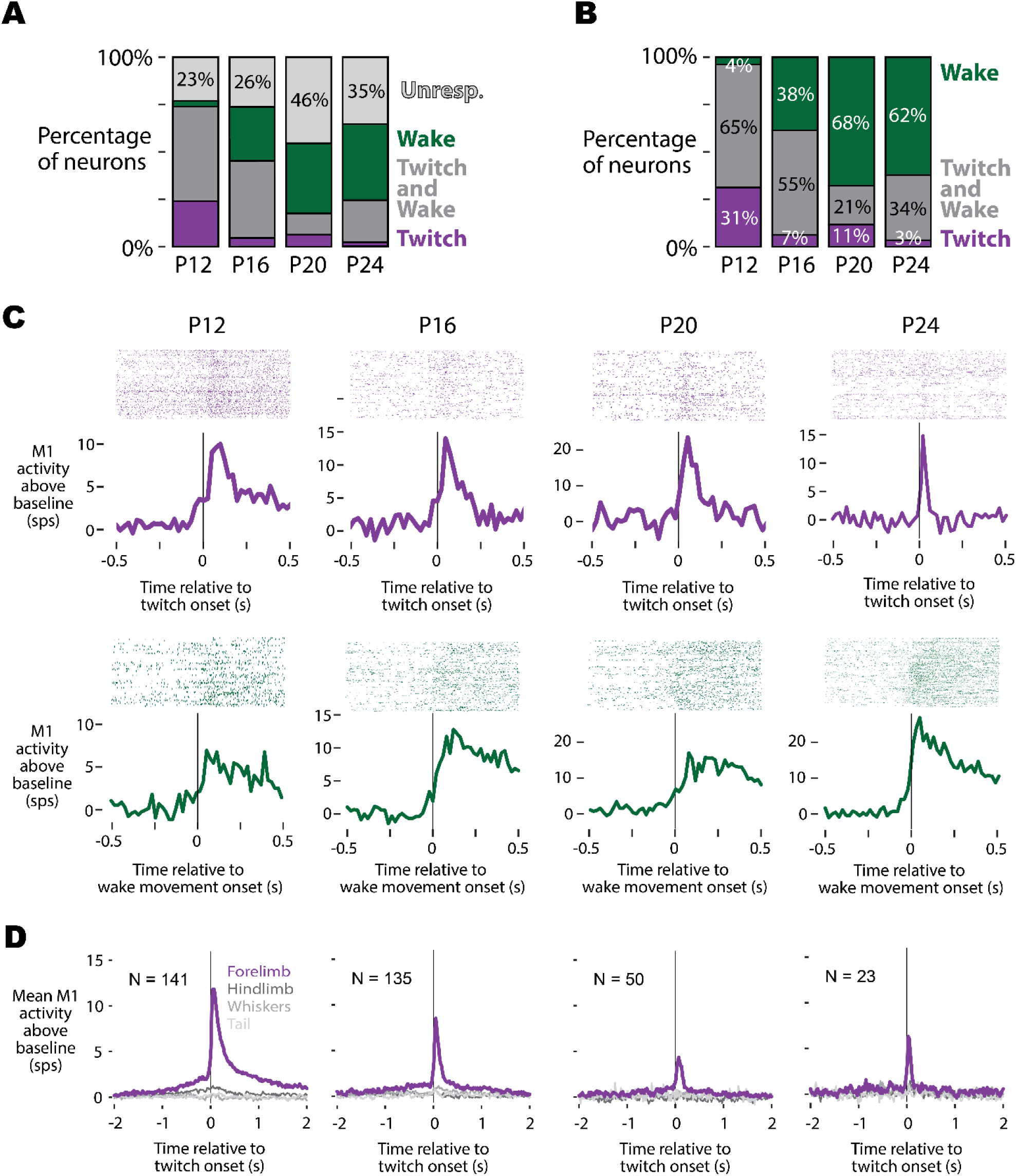
Movement-related activity in M1 neurons through P24. **(A)** Stacked bar plots showing the percentage of M1 neurons at each age (P12, P16, P20, P24) that respond to self-generated movements (i.e., twitches during REM sleep or wake movements during wake). **(B)** Of the movement-responsive neurons identified in **A**, the percentage that respond to twitches (purple), wake movements (green), or both (gray) are shown in stacked bars for each age. **(C)** Representative M1 neurons from each age, illustrating raster plots (top) and perievent histograms (bottom) triggered on twitch (purple) or wake movement onset (green). Activity plots depict firing rate relative to baseline, demonstrating strong but developmentally evolving movement-related activity in M1. **(D)** Mean perievent firing rate of all twitch-responsive M1 neurons at P12 through P24 triggered on isolated twitches of the forelimb, hindlimb, whiskers, and tail. Each panel shows neural activity aligned to twitch onset (time 0). At all ages, neurons in the forelimb region of M1 exclusively respond to forelimb twitches, whereas twitches of other body parts (hindlimb, whiskers, tail) fail to evoke a neural response.

Because twitches are discrete, often involving only a single muscle at a time, we were able to use these movements to examine whether M1 shows somatotopic precision at these ages. We scored twitches of the right forelimb, right hindlimb, whiskers, and tail, enabling us to trigger M1 neural activity on temporally-isolated twitches of all 4 of these body parts (i.e. forelimb twitches that do not occur within 100 ms of twitches of the hindlimb, whiskers, or tail). Our data show that, at all ages examined (P12 to P24), twitch-responsive neurons in the forelimb region of M1 only respond to forelimb twitches (**Figure 2D**). This demonstrates that at least coarse (within a limb) somatotopy is present in M1 by P12.

### M1 responses become faster and more temporally precise with age

We next examined whether the timing and duration of M1 responses becomes more refined with age, a process previously reported in thalamus (Dooley et al., 2021) that is likely necessary for the emergence motor outflow. At a population level, the overall shapes of twitch- and wake-movement-related activity was similar across ages, with twitch-related activity being briefer and more symmetric than wake-movement-related activity. (**Figure 3A, B**). However, twitch-responsive neurons in M1 appeared to show temporal refinement across age. To test this more quantitatively, we performed several kinematic analyses of twitch-related activity. We restricted these kinematic analyses to twitches since these movements are discrete and

**Figure 3.**
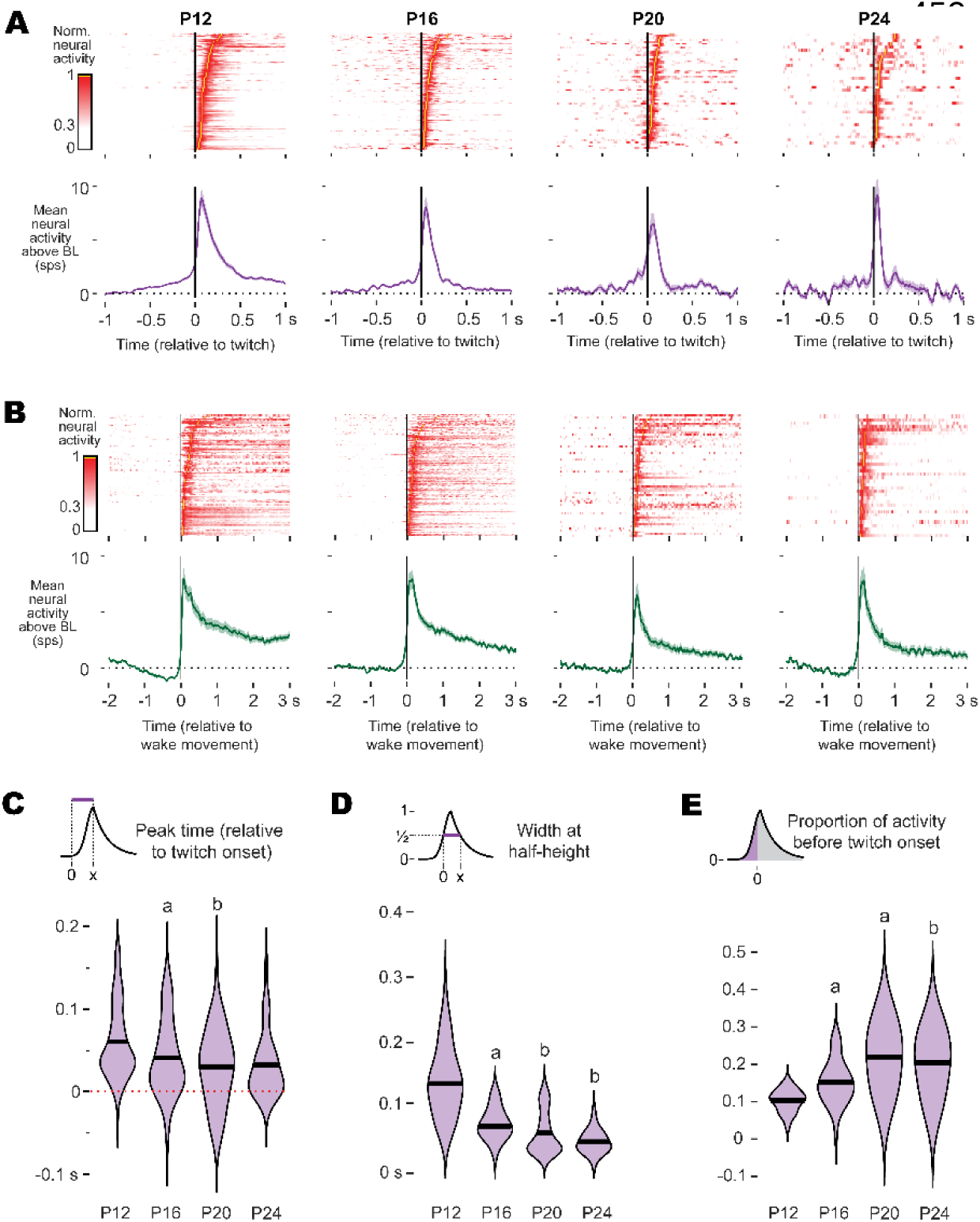
Temporal refinement of twitch- and wake-related M1 activity through P24. **(A)** Top: Heatmaps showing twitch-triggered responses of all twitch-responsive M1 neurons across P12, P16, P20, and P24. Each row corresponds to a single neuron, and color intensity indicates firing rate. Bottom: Mean (±SEM) perievent histograms of these neurons for each age. **(B)** Same as **A**, but for wake-triggered responses of M1 neurons. **(C)** Violin plots of the peak response time (relative to twitch onset) for twitch-responsive M1 neurons by age. The black line represents the mean of the distribution. ‘a’ indicates a significant difference from the preceding age, whereas ‘b’ indicates a significant difference from two ages prior. **(D)** Same as **C**, but for the width at half-height of twitch-triggered responses. **(E)** Same as **C**, but for the proportion of twitch-related activity that precedes movement. Collectively **C-E** reveals progressive temporal sharpening of M1 responses to twitches over the course of development.

### M1 responses become fasterand more temporally precisewith age

We next examined whether thetiming and durationof M1 responses becomes more refinedwith age, aprocesspreviouslyreported in thalamus(Dooley etal., 2021)that is likelynecessaryfor theemergencemotor outflow.At a population level, the overallshapes of twitch-and wake-movement-relatedactivitywassimilar across ages,with twitch-related activitybeing brieferandmore symmetric thanwake-movement-related activity.(**Figure3A, B**). However, twitch-responsive neurons in M1 appeared to show temporalrefinement across age.To test thismore quantitatively,weperformedseveralkinematic analyses oftwitch-related activity. Werestricted these kinematicanalysesto twitchessince thesemovementsare discrete andconsistentacross these ages(**Figure S3B**), whereas wakemovements vary widelywithrespect to boththe timing ofpeakdisplacementand duration (**Figure S3C**), preventingadirectcomparison ofduration and timingfor wake-related activityacross ages.

The time of the peak response of twitch-related activityvariedsignificantly with age (F(3,305)= 7.28; p <0.0001);older animalsshowed earlierpeak activity thanyounger animals (**Figure3C**), such that at P20 and P24, thetiming ofpeak M1 responsesmore closely resembled the timing observed in the ventral lateral nucleus (“motor” thalamus) than the ventral posterior nucleus of the thalamus (see Dooley et al., 2021). In addition, response durations shortened significantly with age, as shown by narrower widths at half-height (F_(3, 209)_ = 60.4; p < 0.0001; **Figure 3D**), indicating that older animals generated briefer, more temporally precise responses to twitches.

To test whether this refinement included emerging signs of premovement activity, we quantified the proportion of spikes occurring before twitch onset (**Figure 3E**). This measure is influenced by the overall duration of responses, which are longer at younger ages, but provides insight into whether twitch-related activity in M1 shows a systematic shift by age. We observed a significant shift from P12 to P16, and then again from P16 to P20, with the activity preceding movement increasing significantly with age (F_(3, 216)_ = 30.5; p < 0.0001), from a mean proportion of 0.10 at P12 to a mean proportion of 0.22 and 0.20 at P20 and P24, respectively. Thus, by P20, about 20% of twitch-related activity in M1 precedes movement onset.

Altogether, we observed an increase in the proportion of twitch-related activity that precedes movement across these ages (see **Figure 3E**), suggesting that by P24, activity in M1 may be beginning to show features necessary of motor output center like RN, particularly an increase in the proportion of premovement spiking. Yet without comparison to a subcortical motor structure that is known to drive infant behaviors, the functional significance of premovement activity in M1 remains unclear. Thus, to better contextualize this transition, we next compared activity in M1 of P24 rats with activity in RN, the midbrain motor nucleus capable of producing movements, including twitches, in early infancy (Williams et al., 2014; Rio-Bermudez et al., 2015).

### Movement-related activity in RN at P24

Prior work demonstrates that RN generates forelimb twitches through at least P12 (Rio-Bermudez et al., 2015; Del Rio-Bermudez et al., 2016), and work on M1 and RN interactions suggests that RN’s role in motor control persists through the developmental emergence of M1 motor control (Williams et al., 2014; Williams and Martin, 2015) if not throughout the lifespan (Gassel et al., 1965). Thus, in the absence of cortical motor control, RN is presumed to be responsible for forelimb movements through P24, although this has yet to be demonstrated. To investigate RN’s role in movement production and compare its activity to M1, we performed dual recordings in M1 and RN of P24 rats (**Figure 4A, B**; N = 7 animals; 112 Neurons in M1; 111 Neurons in RN). In RN, baseline firing rates significantly differed across REM sleep and wake (t_110_ = 8.86, p < 0.0001) with RN firing nearly twice as much during REM sleep (x̂ = 13.1 spikes/s) than during wake (x̂ = 7.2 spikes/s; **Figure 4C**).

**Figure 4.**
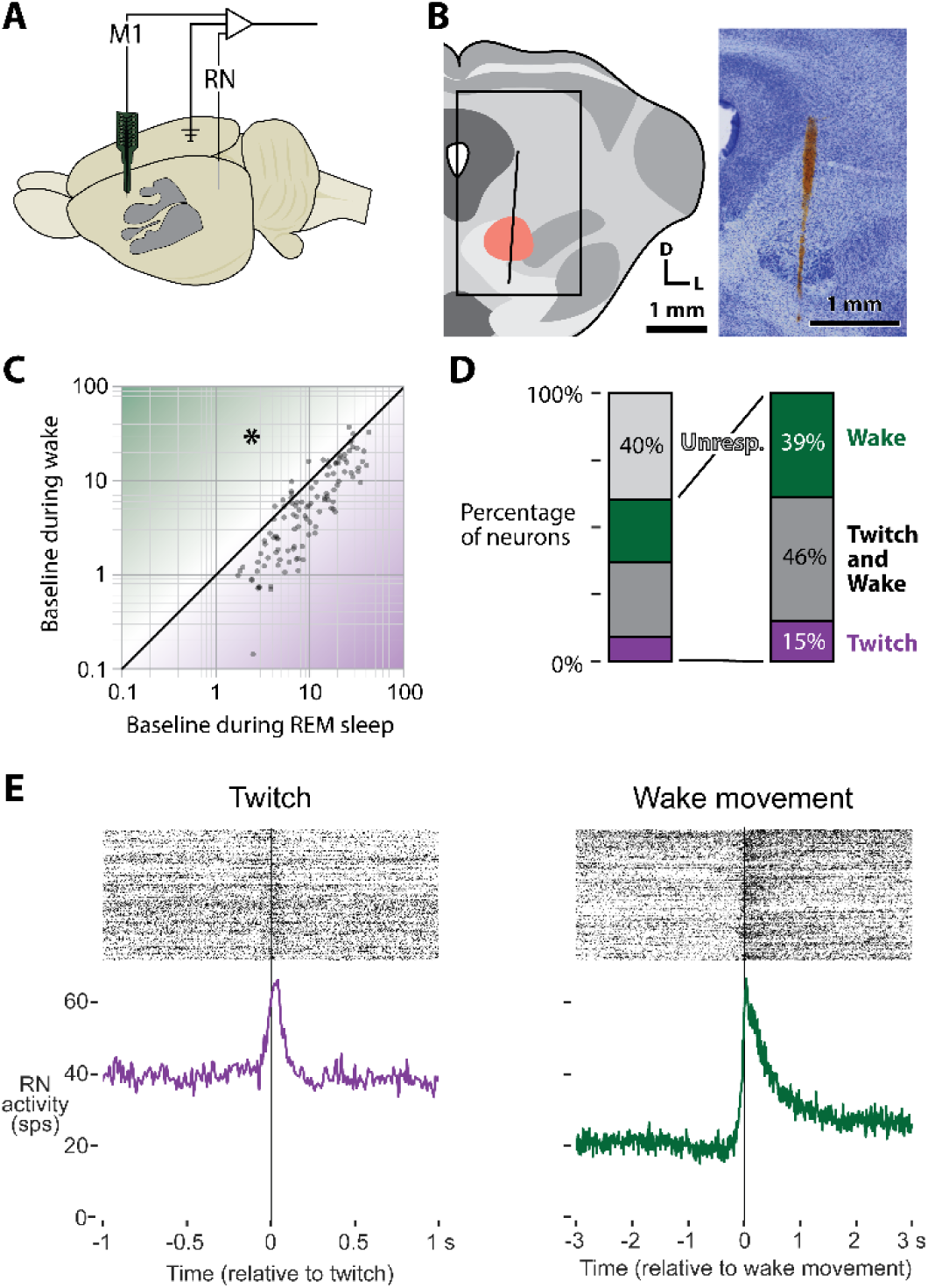
RN recordings and responsiveness to movement at P24. **(A)** Schematic of the rat brain illustrating the dual recording sites in M1 and RN in P24 rats. **(B)** Illustration of a coronal section through the midbrain, showing the RN (red oval) and a representative electrode track for RN recordings. The rectangle on the left indicates the region enlarged on the right, which depicts the Nissl-stained section with the electrode track (red) passing through RN. **(C)** Scatter plot of baseline RN neuronal activity during REM sleep (x-axis) versus wake (y-axis). Most RN units exhibit higher firing rates in REM sleep than in wake, and this difference is significant (asterisk). **(D)** Stacked bar plots showing the proportion of RN neurons responsive to forelimb movements. The subplots on the right illustrate the fraction of those responsive neurons that react to twitches (purple), wake movements (green), or both (gray). This parallels the format of Figures 2A and 2B for M1. **(E)** Representative RN neuron with raster plots (top) and averaged perievent histograms (bottom) for twitch- (left, purple) and wake-movement-triggered (right, green) responses.

A majority of RN neurons from P24 rats showed an increase in activity during forelimb movements (60%; **Figure 4D**). 61% of the forelimb-responsive neurons showed increased activity around twitches, while 85% of forelimb-responsive neurons showed increased activity around wake movements (**Figure 4D**). A representative RN neuron showing twitch- and wake-movement-related activity is shown in **Figure 4E**. Thus, RN maintains strong movement-related activity through P24.

### Movement-related activity in M1 lags behind RN at P24

Having established that RN maintains movement-related activity across sleep and wake through P24, we next asked how its activity compares to M1. The population level activity of P24 rats in M1 and RN are shown for twitches and wake movements in **Figures 5A and 5B**, respectively, with M1 population activity in this second cohort of P24 rats being very consistent with the activity reported earlier (see **Figures 2 and 3**).

**Figure 5.**
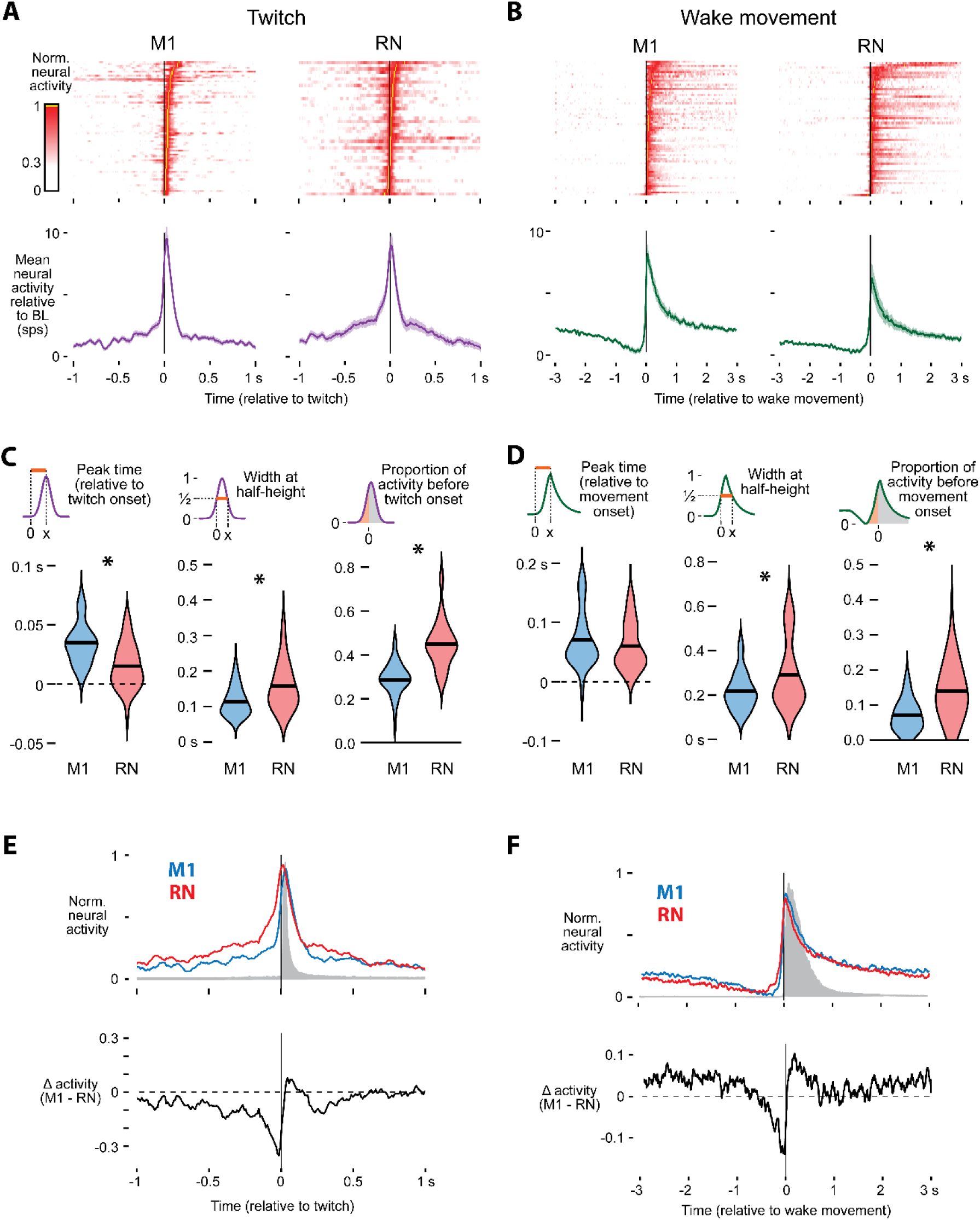
Comparison of twitch- and wake-movement-related activity in M1 and RN at P24. **(A)** Heatmap of mean twitch-related activity of all twitch-responsive M1 (left) and RN (right) neurons recorded during dual M1–RN recordings. **(B)** Same as **A**, but for wake movements. **(C)** Violin plots comparing twitch-responsive neurons in M1 (blue) and RN (red). From left to right: peak response latency, response width at half-height, and proportion of spikes occurring before twitch onset. RN responses peaked significantly earlier, were broader, and contained a greater fraction of premovement activity than M1 responses. Asterisks indicate significant group differences. **(D)** Same as **C**, but for wake movements. **(E)** Top: Average twitch-related activity of M1 (blue) and RN (red) neurons, normalized to 1. The gray shaded region indicates median displacement from the twitch (see **Figure S3**). Bottom: Difference in normalized activity (M1 minus RN) shows twitch-related activity in RN precedes M1 and twitch onset. **(F)** Same as **E**, but for wake movements.

Despite their superficially similar profiles of movement-related activity in M1 and RN, there are several key differences. Compared to twitch-related activity in M1, RN shows earlier peaks (RN: 15 ms, M1: 34 ms; t_98_ = −5.18, p < 0.0001), broader responses (t_102_ = −4.18, p < 0.0001), and a greater proportion of spikes preceding twitch onset (RN: 0.45, M1: 0.29; t_109_ = −8.35, p < 0.0001; **Figure 5C**). Unlike twitches, peak timing did not differ between RN and M1 during wake movements (t_106_ = −1.22, p = 0.23), but RN still showed broader responses (t_110_ = 3.16, p = 0.002) and a greater proportion of premovement activity than M1 (RN: 0.14, M1: 0.07; t_115_ = −5.70, p < 0.0001; **Figure 5D**). Critically, because the recordings in M1 and RN were simultaneous, the same animals movement-related activity was triggered on the same movements. Thus, these differences reflect differences in neural activity, not movement variability. Across both behaviors, activity in RN consistently rose earlier than M1, which can be revealed by subtracting normalized movement-related activity in M1 and RN (**Figure 5E, F**). This is consistent with RN showing stronger premovement spiking and reveals a relative rebound in M1 following movement onset.

In adults, individual neurons in both RN and M1 fire at precise times during the execution of distinct behaviors such as locomotion or reaching (Rho et al., 1999; Hermer-Vazquez et al., 2004; Estebanez et al., 2017; Viaro et al., 2021). In contrast, infant M1 appears to respond broadly and non-selectively, showing activity across many different movements. By comparing movement variability with response variability in M1 and RN, we can test whether either structure begins to show adult-like selectivity during particular wake movements at P24.

### RN neurons are more movement-selective than M1 neurons

When we examined movement-related activity in RN at P24, we found diverse and heterogeneous responses, particularly during wake movements, consistent with a role in generating distinct behaviors. We often observed individual neurons in RN showing increased activity for some wake movements and suppression for others (**Figure 6A, top**), a pattern that was not seen in M1. This difference in the variability of movement-related activity was confirmed quantitatively: We found that wake-movement-related activity showed significantly higher response variability in RN than in M1, as indicated by a higher coefficient of variation in RN (t_211_ = −4.93, p < 0.0001; **Figure 6B**).

**Figure 6.**
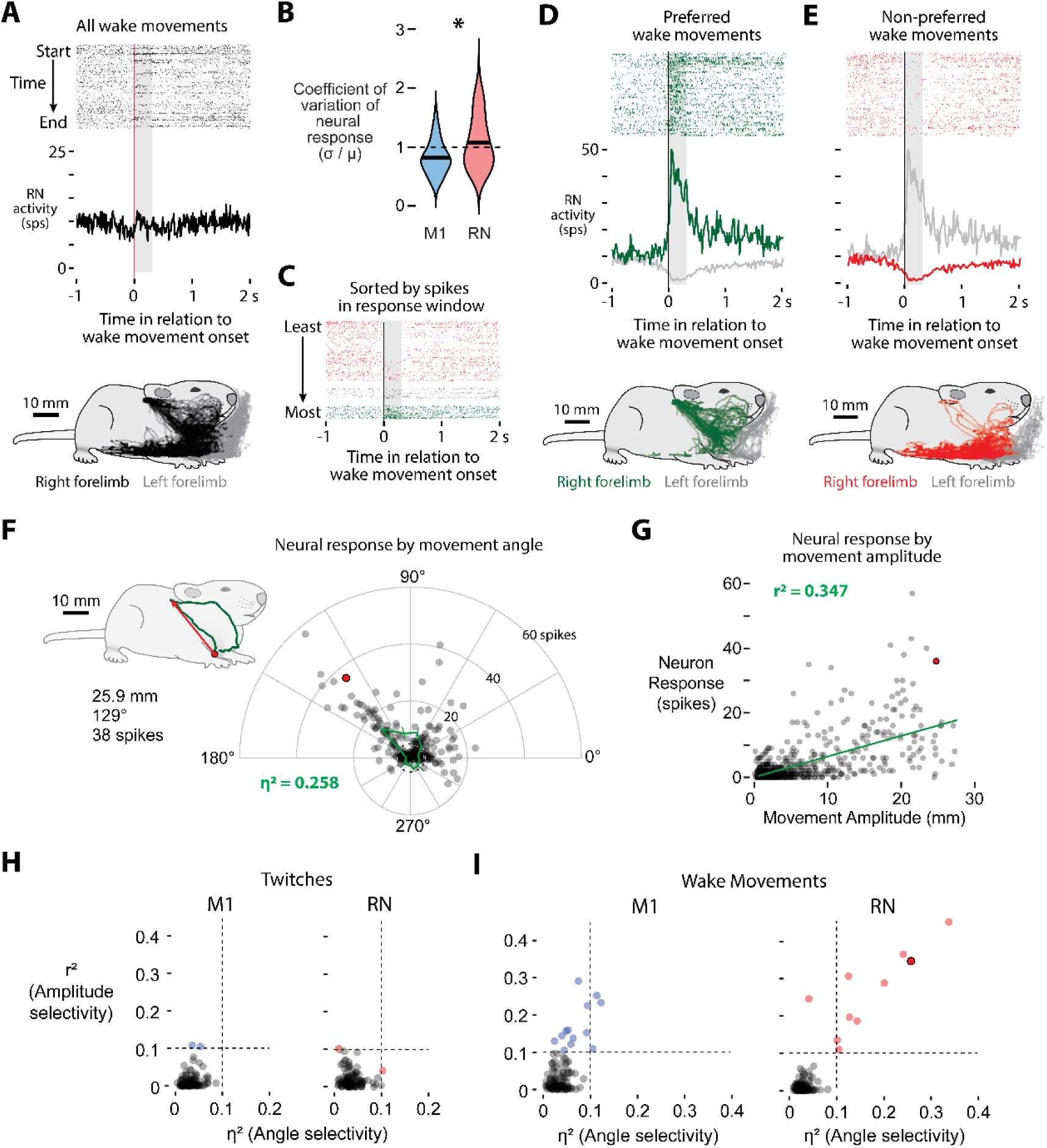
RN neurons are more selective to specific wake movement than M1 neurons. **(A)** Top: Raster and perievent histogram for all wake movements of a representative RN neuron that responds to only a subset of wake movements. Bottom: Location of the right and left forelimbs (black and grey, respectively) for all scored wake movements. **(B)** Violin plots of the coefficient of variation of neural responses to wake movements for all M1 and RN neurons. Wake movement activity in RN is significantly more variable than M1. **(C)** The raster plot shown in **A**, sorted by the number of spikes in the response window. **(D)** Top: Raster and perievent histogram for the preferred wake movements. Bottom: Location of the right and left forelimbs (green and grey, respectively) for wake movements accompanied by a neural response. **(E)** Same as in **D**, but for non-preferred wake movements. **(F)** Polar plot for a representative RN neuron showing spike counts as a function of wake-movement direction (12 bins). The inset traces one example movement: the right forelimb starts at the red circle and reaches a maximum displacement of 25.9 mm at 129° (red arrow). This movement is highlighted with a red dot here and in **G**. Direction accounts for 25.8% of the neuron’s response variance (η² = 0.258). **(G)** The same neuron’s firing versus movement amplitude. The green line is the linear fit (r² = 0.347). **(H)** For twitches, neither movement angle (η²) nor amplitude (r²) explains much response variance in M1 (left) or RN (right); only two neurons in each region exceed the 0.1 threshold (dashed lines). **(I)** For wake movements, a larger subset of M1 neurons shows amplitude selectivity (elevated r²), whereas RN neurons more often exhibit combined angle- and amplitude-tuning (both η² and r² > 0.1). The red-outlined point marks the example neuron from **A**–**G**.

Next, we wanted to see if this neural response variability in RN tracks with movement variability. We applied DeepLabCut to trace the limb position of the right- and left-forelimbs at every frame (**Figure S4A, B**), revealing the diverse and highly variable kinematics of wake behaviors (see **Figure 6A, bottom**). If individual neurons drive individual single behaviors, in a naturally behaving animal, specific neurons should fire when an animal performs a specific movement.

Taking the neuron in **Figure 6A**, we categorized each wake movement by the number of RN spikes within a 300 ms response window: events with more than 50% fewer spikes than expected in this window from baseline activity alone (2 or fewer spikes) were classified as “non-preferred,” whereas those with 50% more spikes than expected from baseline activity (6 or more spikes) were classified as “preferred” (**Figure 6C**). This yielded two sharply contrasting perievent histograms, one broadly excitatory and the other broadly inhibitory throughout the movement (**Figure 6D, E**). Strikingly, the ‘preferred’ category isolated a specific, stereotyped behavior—right forelimb movements to the ear and vibrissae (**Figure 6D, bottom**)—which video analysis confirmed to be predominantly bilateral grooming (χ^2^ = 37.9, N = 454, p < 0.001). In contrast, the “non-preferred” category comprised the rest of the wake repertoire, including postural adjustments and locomotion (**Figure 6E, bottom**).

Thus, wake-movement-related activity in the RN is not always uniform; the same neuron switches from excitation to inhibition depending on the movement. Similar mixed responses have been shown in M1 of adult rodents (Peters et al., 2017), hinting that distinct movement categories differentially recruit these circuits. Motivated by this, we performed a kinematic analysis of forelimb movements to test whether RN and M1 neurons respond to movements with specific amplitudes or direction. To do so, we used limb position data to quantify the angle and amplitude of both twitches and wake movements and related these measures to neural activity on a movement-by-movement basis.

Not surprisingly, wake movements were much larger in amplitude than twitches, with wake movements averaging 6.95 mm and twitches averaging 0.96 mm (**Figure S4C**). The amplitude and angle of all wake movements and twitches can be seen in **Figures S4D and E**, respectively (N = 4365 wake movements, N = 3766 twitches; pooled across 7 animals). A majority of right forelimb movements were forwards or backwards from a resting position (0° and 180°), which during wake consists largely of locomotion and postural control movements. Because the pup’s limbs are resting on the surface of the Mobile HomeCage, very few movements were in the downward direction (270°), however a subset of movements were in the upwards direction, (90°) towards the face. Video analysis confirmed that these movements were predominantly facial grooming. Twitches were more uniformly distributed than wake movements, although still biased to be forwards or backwards.

After determining twitch and wake movement trajectories, we next established whether movement direction and movement amplitude explain variability of M1 or RN neural responses. For each neuron, we quantified the number of spikes within the “response window” of each movement (−70 to 70 ms for twitches, and −100 to 500 ms for wake movements). We then calculated how much of the response variability is explained by movement amplitude (via a regression) or movement direction (quantified by grouping movements in 12 equally sized directional bins; see **Figures S4D, E** and **6F**).

The response variability explained by movement angle and movement amplitude of the neuron in **Figure 6A** is shown in **Figures 6F** and **6G**, respectively. A single representative grooming movement (**Figure 6F inset**) is illustrated with the red dot, while all other wake movements are illustrated with a gray dot. This neuron showed a strong response to movements towards the face and ears (60° to 150°) and a weaker than average response to movements in other directions, such that movement direction explained 25.8% of neural response variability. Likewise, movement amplitude was positively correlated with neural response, explaining 34.7% of response variability. For this neuron, twitch-related activity did not meaningfully explain response variability, with movement direction and amplitude explaining 4.5% and 0.1% of response variability, respectively (data not shown).

Across the population, limb kinematics did not predict twitch-evoked firing in either M1 or RN (**Figure 6H**). But during wake movements, this kinematic analysis revealed a critical difference: although only a minority of neurons were selective (13 of 112 in M1, 10 of 111 in RN), RN neurons showed a strikingly different profile. In RN, nearly every selective neuron (9 of 10; see **Figure 6I**) showed selectivity for both amplitude and direction, clearly encoding movements to specific locations. In contrast, nearly all selective M1 neurons (10 of 13) showed selectivity only for amplitude. And although three neurons surpassed the threshold for direction selectivity, their direction tuning was relatively weak, barely exceeding our threshold (η² = 0.1). Thus, the coupling of amplitude and direction selectivity in RN contrasts with the amplitude-based selectivity of M1, consistent with RN neurons participating more directly with specific motor commands and M1 neurons representing a broader mix of signals, including predicted and actual sensory feedback.

## Discussion

By recording neural activity in M1 in P12 and P24 rats as they cycled between sleep and wake (**Figures 1 and S1**), we showed that a subset of neurons in M1 show twitch- and wake-movement-related activity through P24 (**Figure 2**). Because twitches are discrete and stereotyped, they provide a consistent measure of movement-related activity across ages. Exploiting this, we found that by P12, M1 is already somatotopically organized, and its twitch-related responses become progressively faster and sharper across age, including an increase in premovement activity (**Figure 3**). However, simultaneous recordings in M1 and RN reveal a functional difference: in P24 pups, neurons in RN not only fire earlier than M1 for both twitches and wake movements (**Figures 4 and 5**) but also display far stronger tuning to the direction and amplitude of individual wake movements (**Figure 6**). Thus, while M1 is refining its sensory representation of movement, RN activity is consistent with its role as the principal driver of limb movements, with individual RN neurons firing selectively during specific wake movements. By providing these structured motor signals, neurons in RN supply the movement-related activity that shapes M1 through experience-dependent plasticity.

### The same movements drive different patterns of M1 activity across development

Although infant M1 is not yet producing movement, it is not silent. M1 undergoes several functional transformations prior to assuming its adult role producing behaviors. From near birth through P10, activity in M1 is strongly patterned by twitches, whereas wake-movement-related activity is inhibited, mirroring the broader sensorimotor system’s early reliance on reafferent signals from twitches (Khazipov et al., 2004; Tiriac et al., 2014; Tiriac and Blumberg, 2016; Dooley et al., 2020). From P10 to P12, functional activity in M1 undergoes several changes. The first change is stopping the inhibition of wake-movement-related activity; a transition that also occurs in the developing somatosensory and visual cortices around this age (Dooley and Blumberg, 2018; Murata and Colonnese, 2018). The second change is in the types of somatosensory inputs that M1 receives. Prior to P12, M1 receives both tactile and proprioceptive inputs from thalamus, in parallel with somatosensory cortex. From P12 onward, M1 only receives proprioceptive inputs from thalamus, becoming reliant on somatosensory cortex for tactile information (Gómez et al., 2021), again mirroring a similar change in parallel to serial processing seen in the developing visual system (Murakami et al., 2022). Finally, population-level activity in M1 goes from bursty to continuous while neurons rapidly decorrelate their sensory responses (Glanz et al., 2021). From the present investigation, we also now know that by P12, M1 shows somatotopic refinement, with the forelimb region of M1 only responding to forelimb twitches (see **Figure 2D**).

The present investigation focused on movement-related activity in M1 following the P12 transformation. Prior work on somatosensory and motor thalamus (the ventral posterior and ventral lateral nuclei, respectively) identified P12 to P20 as a period of temporal refinement of movement-related activity (Dooley et al., 2021). Because motor thalamus is the primary thalamic input to M1, it is not surprising that we observed similar temporal refinement in M1 across this developmental window. In addition, by P20, activity in motor thalamus no longer reflects sensory inputs, but instead reflects a cerebellar-dependent internal model of movement. In the present investigation, the mean latency to peak twitch-related activity in M1 at P20 and P24 is approximately 30 ms (see **Figure 3C**)—the same as previously reported in motor thalamus and equivalent to the peak displacement from twitches (**Figure S3**). Thus, by P20, movement-related activity in M1 has likely undergone another transformation, representing a cerebellar-dependent internal model of movement.

### Twitches continue to drive activity in M1 through P24

Prior work at earlier ages demonstrated that twitch-related activity in sensorimotor cortex is quite dynamic across early development (Dooley and Blumberg, 2018; Dooley et al., 2021). Whereas limited evidence in adult animals suggests that REM sleep twitches may drive activity in thalamus (Boscher et al., 2024) and M1 (Marchiafava and Pompeiano, 1964; Eckert et al., 2020), the apparent reduction in the proportion of twitch-responsive neurons in infant sensorimotor cortex (Dooley and Blumberg, 2018; Domínguez et al., 2021; Glanz et al., 2021) has led to the suggestion that the twitch-responsiveness seen in early infancy may be developmentally transient. In line with this notion, our findings reveal that the proportion of twitch-responsive neurons in M1 does decrease from 74% of neurons at P12 to 24% of neurons at P24. Critically, however, a meaningful portion of M1 neurons were still twitch-responsive through P24. This transition, in which M1 neurons shift from responding to twitches to predominantly responding to wake movements, may reflect an expanding range of preferred stimuli as the cortex matures. Indeed, a similar progression—from simple to more complex motor maps—has been documented previously, albeit in rats older than the present investigation (Singleton et al., 2021).

This persistence of twitch-related activity in M1 through P24 suggests that twitch-related activity can continue to play an important role in refining cortical circuits, likely alongside the role of wake-movement-related activity. At this age, neural activity in M1 is critical to the development of cortical motor control (Chakrabarty and Martin, 2005; Williams and Martin, 2015). For the neurons that remain twitch-responsive, reafference from twitchces is an important piece of their overall pattern of neural activity, shaping M1’s circuitry in preparation for its motor functions. These findings keep open the possibility that twitch-related activity continues to shape cortical circuitry through P24, if not longer (Marchiafava and Pompeiano, 1964), underscoring the significance of sleep and twitches in facilitating the final stages of M1’s developmental transition toward adult-like motor control.

### Developmental transition towards motor control

Our dual-site recordings sharpen the contrast between an established motor center (RN) and the still-maturing M1. In P24 pups, RN spiking not only precedes twitches and wake movements (see **Figure 5**) but is tuned to the kinematics of individual wake behaviors. RN neurons fire preferentially for specific forelimb movements and scale their response with movement amplitude (see **Figure 6**). By comparison, activity in M1 lags movement onset (**Figures 3 and 5**) and shows little to no selectivity for movement direction (**Figure 6I**). This is in contrast to M1 in adult rats, which does appear to show selectivity for movement direction (Bonazzi et al., 2013). Thus, whereas RN already shows the movement-specific coding expected of a structure that generates specific behaviors, M1 activity during wake remains dominated by broad responses only scaled by amplitude.

This dissociation implies that the emergence of cortical motor control will be accompanied by a qualitative shift in M1 coding. As corticospinal projections strengthen after P24 and ICMS-evoked movements first appear around P25 (Young et al., 2012; Singleton et al., 2021), we would predict that M1 neurons will begin to display the same relationship between direction- and amplitude-tuning that we observed in RN at P24. In other words, the movement-specific responses that we found in RN should migrate cortically as M1 begins driving behavior. Indeed, a prior study in adult rats shows that both M1 and RN have populations of neurons that show precise responses during specific movements (Hermer-Vazquez et al., 2004). Such a transition would parallel the progressive refinement seen here and in prior work in the timing of M1’s twitch-related activity (Dooley and Blumberg, 2018).

As for when M1 is fully developed, M1’s forelimb motor map appears “adult-like” at P60 (Young et al., 2012; Singleton et al., 2021). However, these motor map milestones are subject to debate regarding what exactly they represent functionally, as some evidence suggests that the formation and reorganization of motor maps lags behind changes of the underlying circuitry (Kleim et al., 2004; Peters et al., 2017). Accordingly, it is plausible that discrete, behaviorally meaningful motor outputs from M1 begin earlier than can be detected via ICMS, and that the progressive refinement we observe in twitch-related M1 activity serves as a key developmental step leading to cortical motor control.

Taken together, our findings support a dynamic, two-stage model. Through P24, neurons in RN initiate specific limb movements (as described by their direction and amplitude), whereas neurons in M1 reflect predicted and actual sensory feedback. As postnatal development proceeds, we expect that M1 will progressively adopt more RN-like movement selectivity. This interplay between M1 and RN likely reflects both the gradual maturation of corticospinal projections and the essential role of early reafferent signals in sculpting motor circuitry (Martin, 2005). Indeed, the incremental “pre-movement” activity we observe in M1 at P20 and P24 may be the first glimpse of its ability to shape behavior. Continuous sensorimotor feedback from both twitches and wake movements likely fuels this process, ensuring that the cortex inherits a well-calibrated, kinematically precise control scheme from brainstem motor centers, rather than creating one de novo.

## Acknowledgments

We thank Mark Blumberg, Greta Sokoloff, and Scott Pluta for feedback on earlier versions of this manuscript. This research was supported by the Sleep Research Society Foundation’s Career Development Award to J.C.D.

## Author contributions

J.C.D. designed research; J.C.D. performed research; M.R.R., N.J.S., and J.C.D. data curation; M.R.R., N.J.S., and J.C.D. analyzed data; M.R.R., N.J.S., and J.C.D. visualization; J.C.D writing – original draft; M.R.R., N.J.S., and J.C.D. writing – review & editing; J.C.D. supervision; J.C.D. resources

## Supplemental Figures

**Figure S1.**
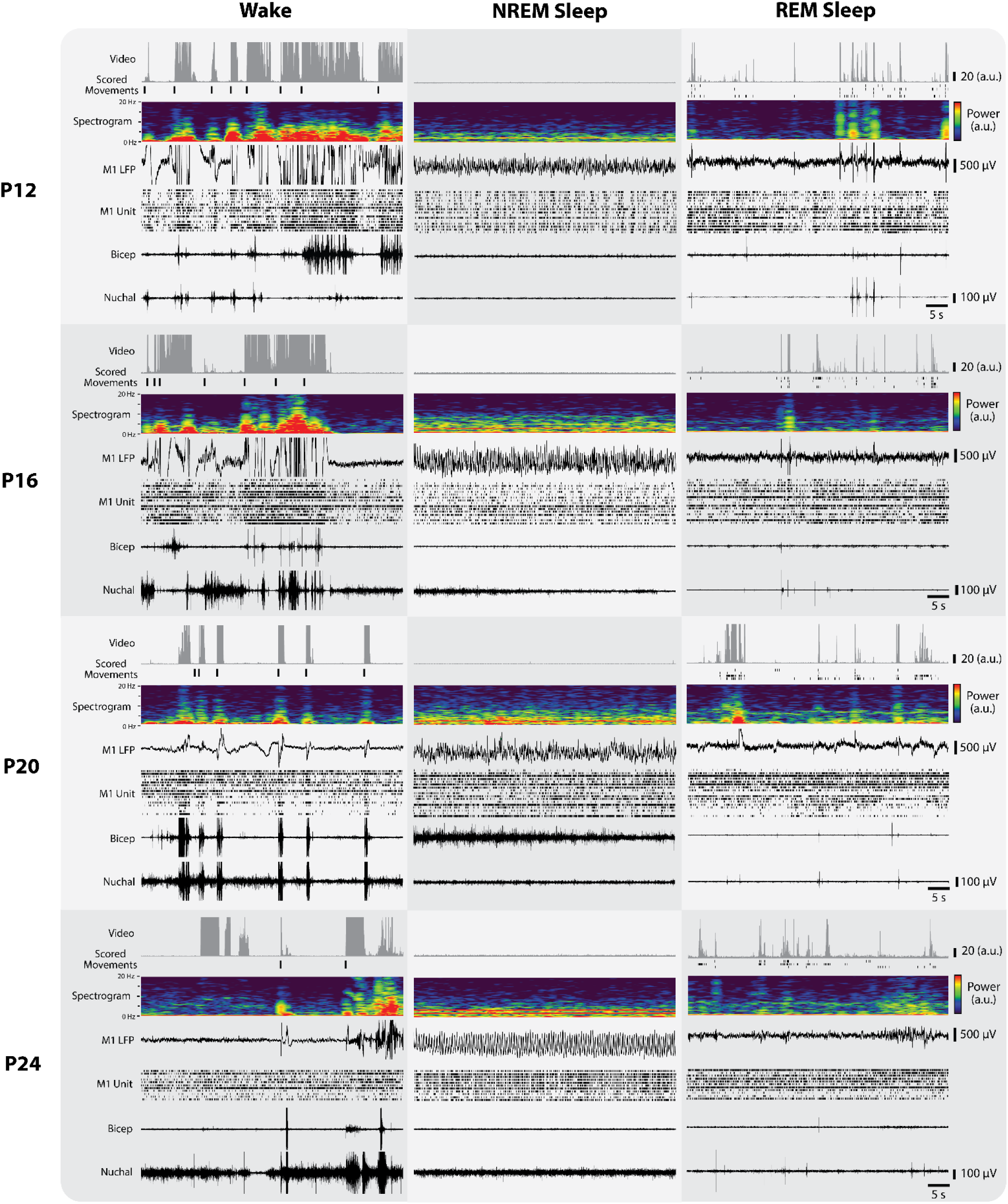
Behavioral and electrophysiological data across wake, NREM sleep, and REM sleep. For each age (P12, P16, P20, P24), a 50-s segment of continuous data is shown for wake, NREM sleep, and REM sleep. Top to bottom: ROI-based movement analysis of the entire rat (as detected by pixel-based motion detection), scored movements (wake movements during wake, twitches during REM sleep), M1 LFP, single-unit activity in M1 (each row corresponds to a separate unit), and biceps and nuchal EMG signals. Rows of scored twitches are forelimb, hindlimb, whisker, and tail, from top to bottom. These representative examples demonstrate distinct patterns of behavioral, neural, and muscular activity at each developmental age.

**Figure S2.**
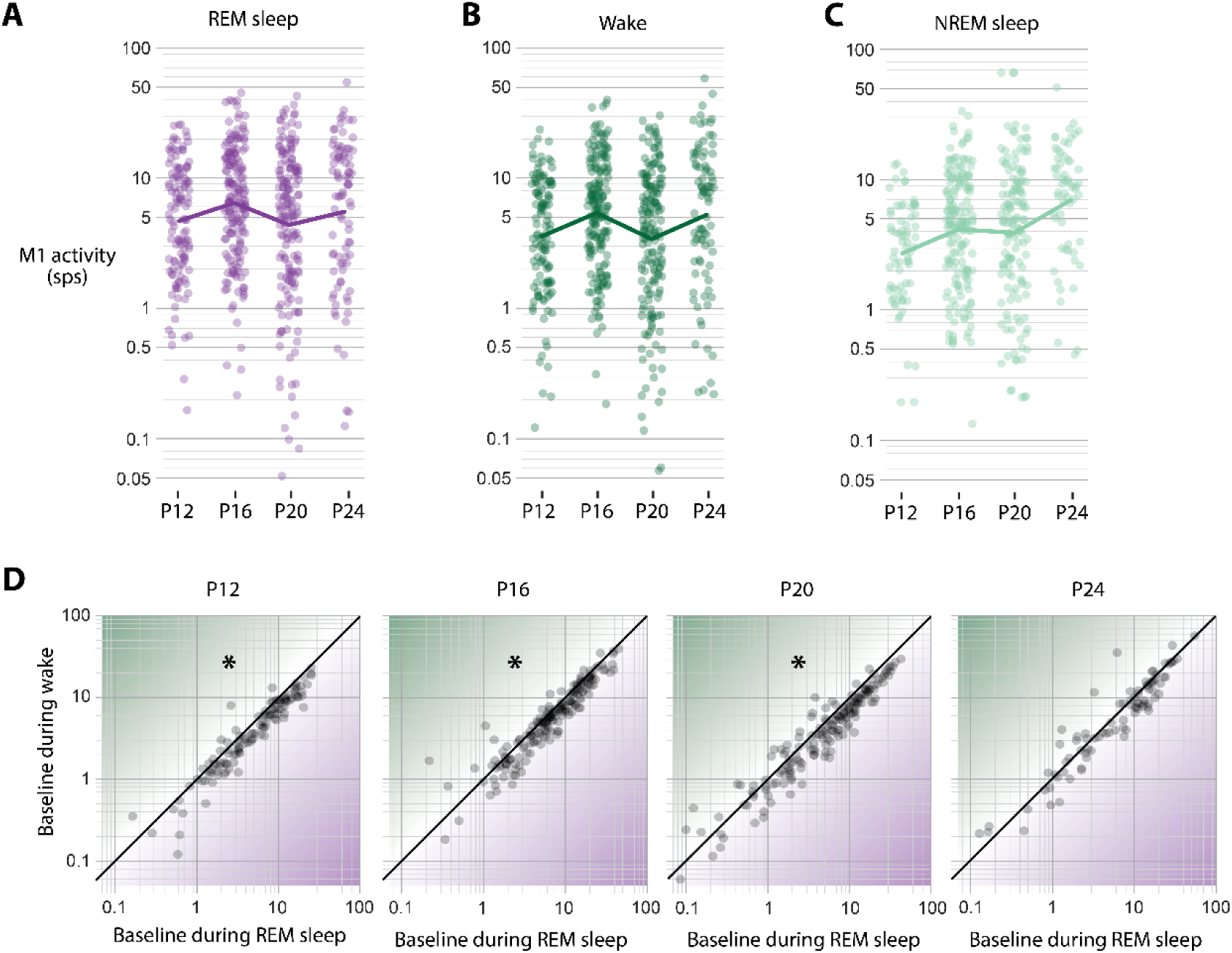
Developmental changes in M1 firing rates across wake and sleep states. **(A)** Log-normalized M1 firing rates for individual neurons (purple dots) during wake at each age (P12, P16, P20, P24). The solid purple line represents the mean log firing rate during wake at each age. **(B)** Same as in (A), but for REM sleep. **(C)** Same as in (A), but for NREM sleep. **(D)** Scatter plot of baseline M1 firing rates for each age during REM sleep (x-axis) versus wake (y-axis). At earlier ages, neurons exhibit significantly elevated activity during REM sleep, whereas this difference diminishes by P24.

**Figure S3.**
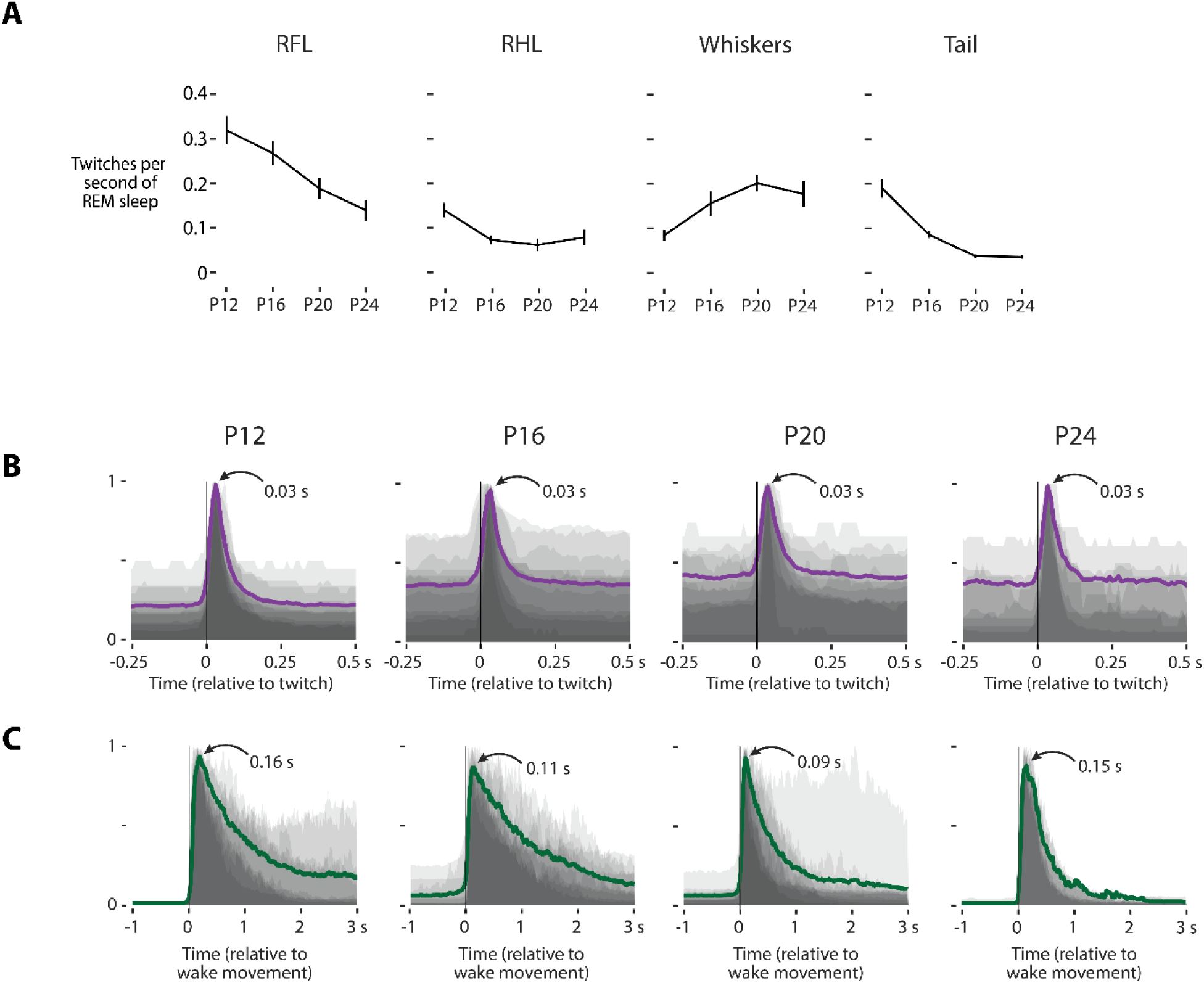
Twitch rates and movement duration by age. **(A)** Mean (±SEM) twitch rates (in twitches per second of REM sleep) for each of the four body parts (forelimb, hindlimb, whiskers, tail) across the four ages (P12, P16, P20, P24). **(B)** Mean displacement produced during right forelimb twitches by age. Each shaded gray region denotes the normalized median displacement produced by that movement for an individual pup. The purple line is the mean across all pups for that age. Across all ages, the mean peak displacement for twitches occurs 0.03 s (or three frames) following twitch onset, demonstrating the duration of twitches is consistent from P12 to P24. **(C)** Same as B, but for wake movements. Unlike twitches, wake movements show variability in the overall mean peak duration and movement duration across ages.

**Figure S4.**
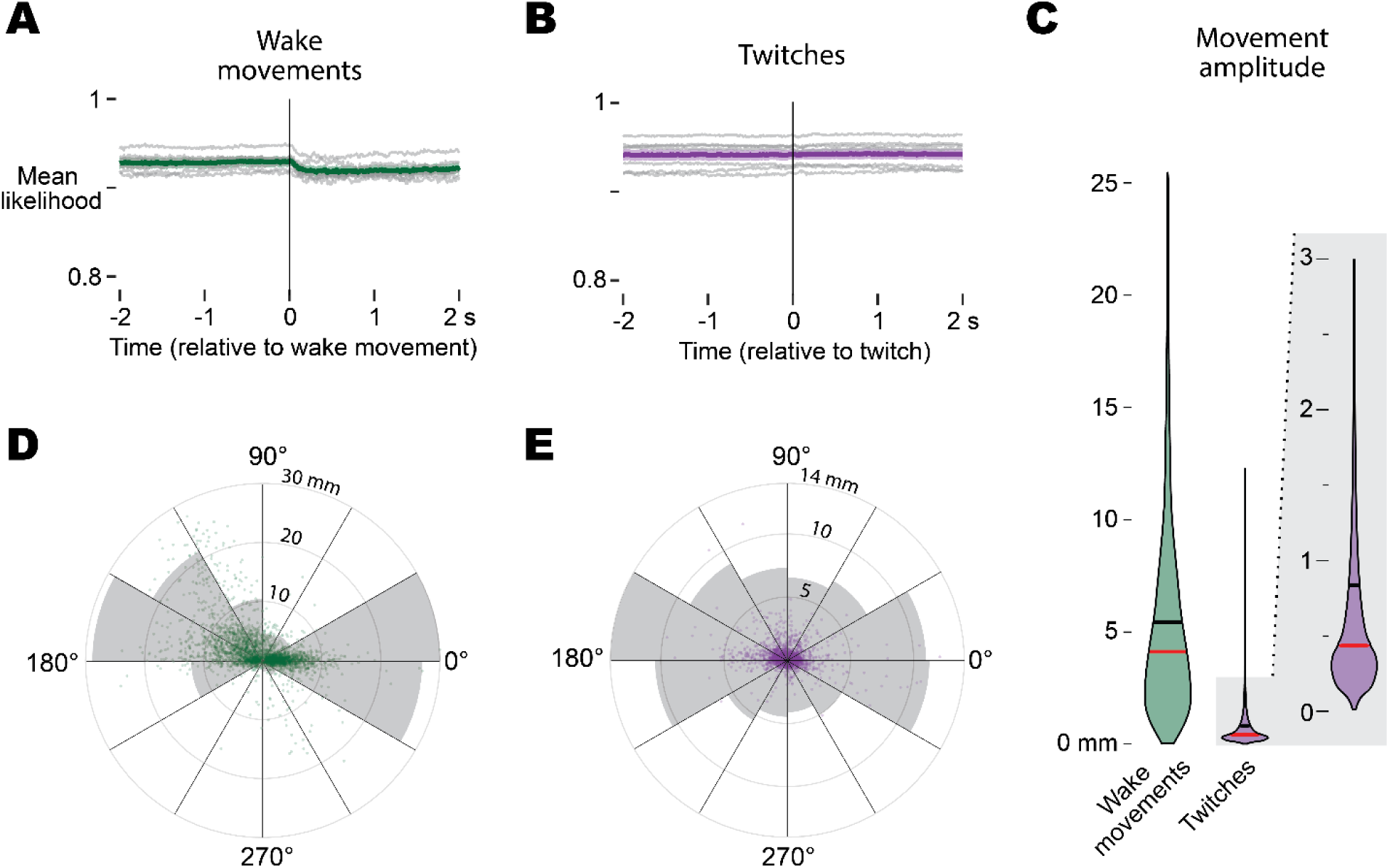
DeepLabCut kinematic summary of wake movements and twitches. **(A)** Mean tracking confidence (likelihood) for right forelimb markers during wake movements. Group average ± SEM is plotted in green; thin grey lines show individual animals (N = 7). **(B)** Same as (A) for REM-sleep twitches, plotted in purple. **(C)** Violin plot comparing peak displacement (movement amplitude) for all wake movements (green) and twitches (purple) pooled across animals (wake n = 4,365; twitch n = 3,766). The black line is the mean and the red line is the median. **(D)** Polar scatter plot of every wake movement: angle denotes movement direction and radial distance indicates amplitude. The grey filled areas are a histogram illustrating the proportion of movements in each directional bin. **(E)** Polar scatter plot of every twitch, formatted as in (D) but shown in purple, highlighting their smaller amplitudes and more evenly-distributed direction.

