## Supplemental Figures for "Subcortically generated movements activate motor cortex during sleep and wake in rats through postnatal day 24"

1010 **Supplemental Figures:**

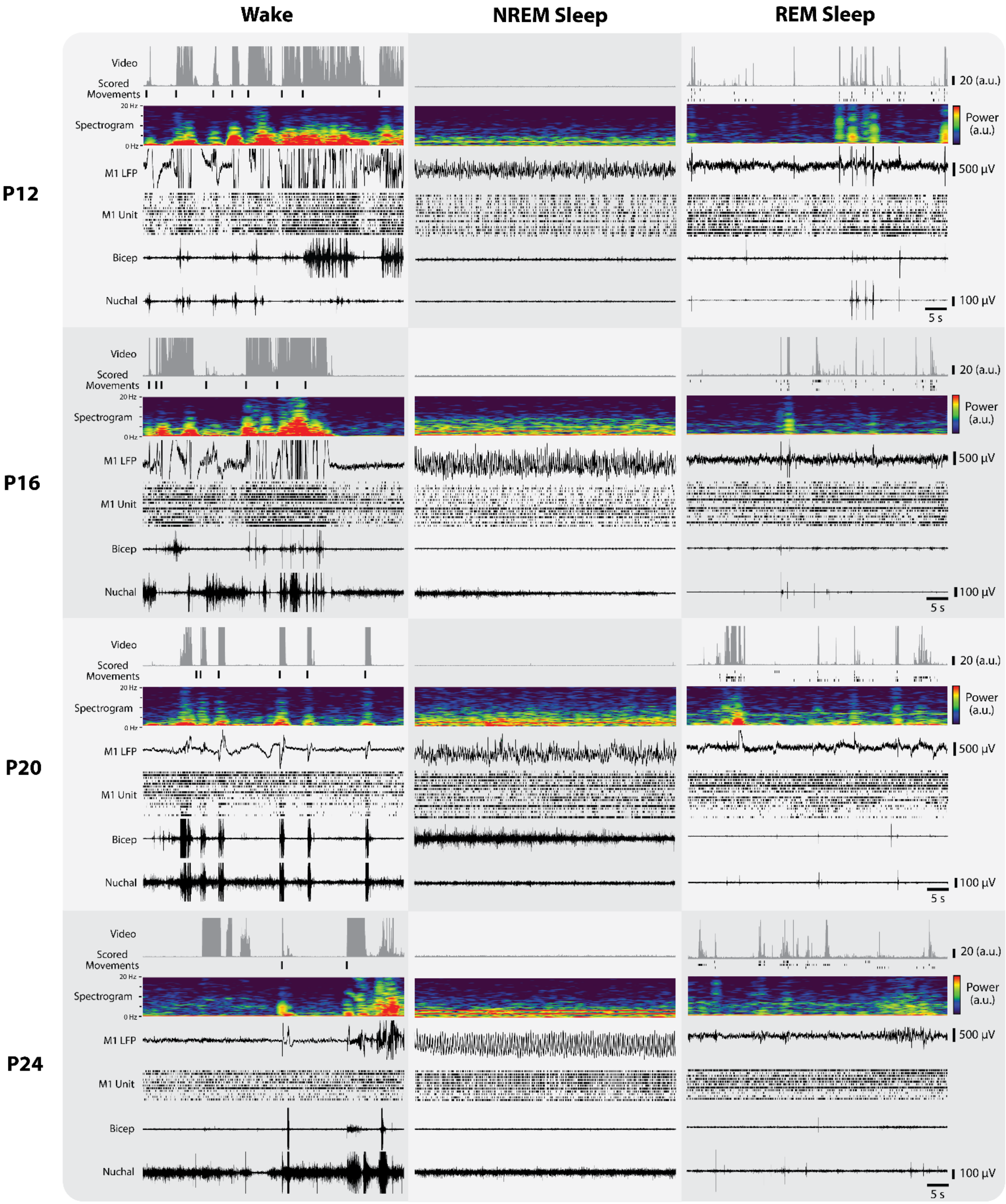


1011

**Figure S1. Behavioral and electrophysiological data across wake, NREM sleep, and REM sleep**

For each age (P12, P16, P20, P24), a 50-s segment of continuous data is shown for wake, NREM sleep, and REM sleep. Top to bottom: ROI-based movement analysis of the entire rat (as detected by pixel-based motion detection), scored movements (wake movements during wake, twitches during REM sleep), M1 LFP, single-unit activity in M1 (each row corresponds to a separate unit), and biceps and nuchal EMG signals. Rows of scored twitches are forelimb, hindlimb, whisker, and tail, from top to bottom. These representative examples demonstrate distinct patterns of behavioral, neural, and muscular activity at each developmental age.

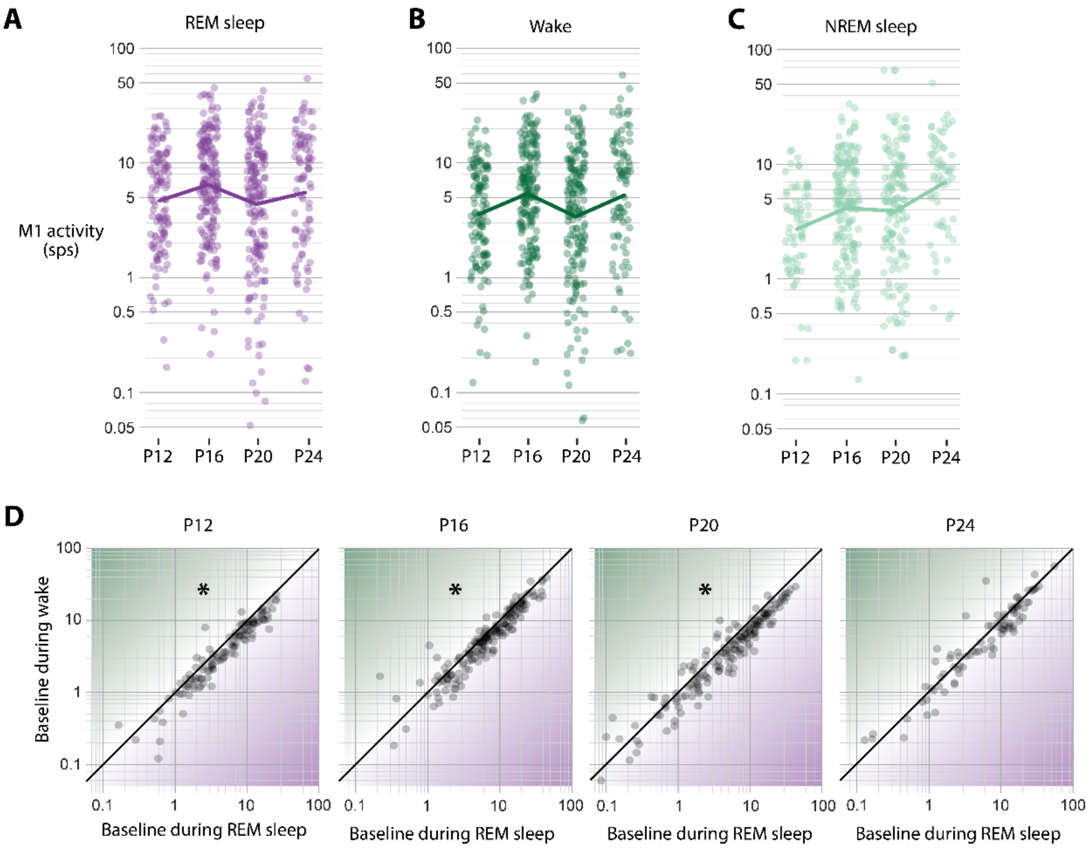


**Figure S2. Developmental changes in M1 firing rates across wake and sleep states**

**(A)** Log-normalized M1 firing rates for individual neurons (purple dots) during wake at each age (P12, P16, P20, P24). The solid purple line represents the mean log firing rate during wake at each age.

**(B)** Same as in (A), but for REM sleep.

**(C)** Same as in (A), but for NREM sleep.

**(D)** Scatter plot of baseline M1 firing rates for each age during REM sleep (x-axis) versus wake (y-axis). At earlier ages, neurons exhibit significantly elevated activity during REM sleep, whereas this difference diminishes by P24.

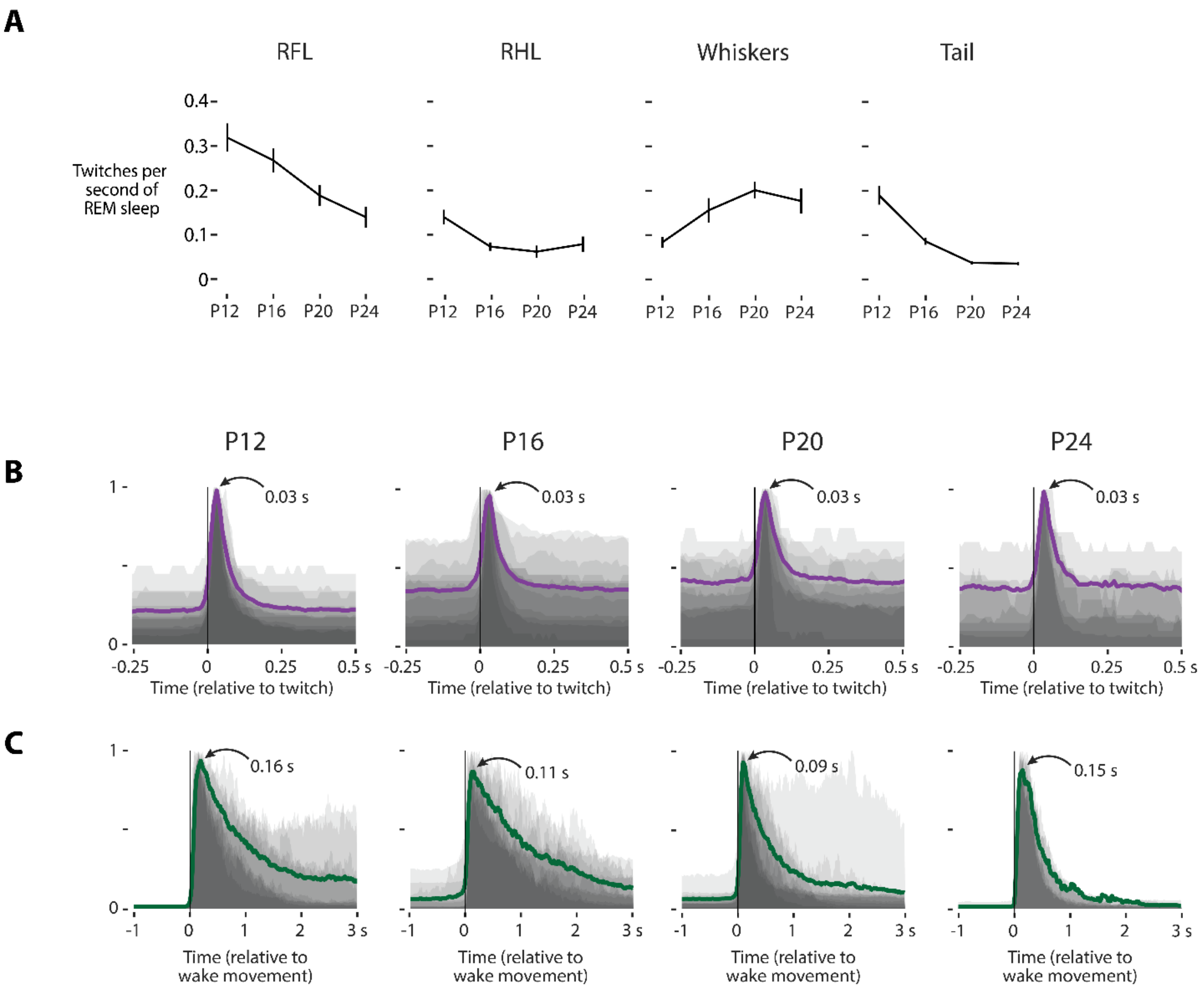


**Figure S3. Twitch rates and movement duration by age.**
**(A)** Mean (±SEM) twitch rates (in twitches per second of REM sleep) for each of the four body parts (forelimb, hindlimb, whiskers, tail) across the four ages (P12, P16, P20, P24).
**(B)** Mean displacement produced during right forelimb twitches by age. Each shaded gray region denotes the normalized median displacement produced by that movement for an individual pup. The purple line is the mean across all pups for that age. Across all ages, the mean peak displacement for twitches occurs 0.03 s (or three frames) following twitch onset, demonstrating the duration of twitches is consistent from P12 to P24.
**(C)** Same as B, but for wake movements. Unlike twitches, wake movements show variability in the overall mean peak duration and movement duration across ages.

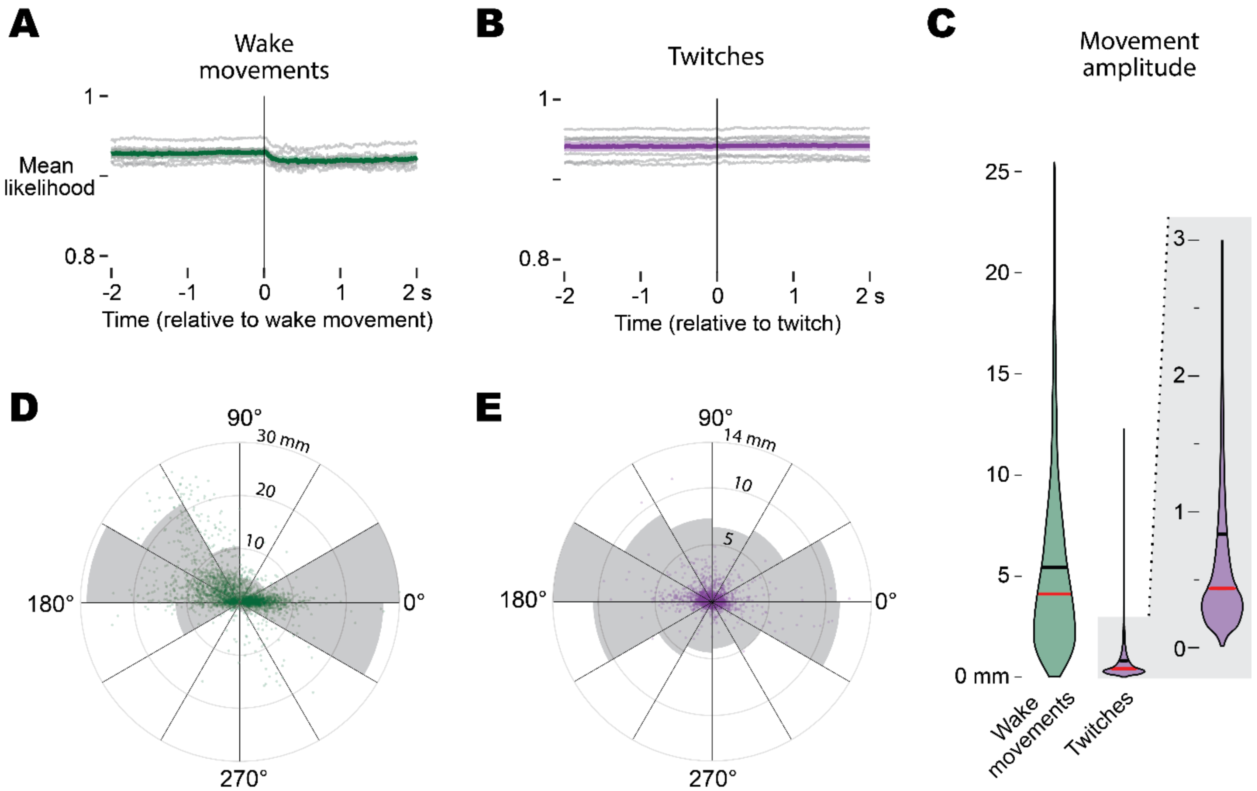


**Figure S4. DeepLabCut kinematic summary of wake movements and twitches**

**(A)** Mean tracking confidence (likelihood) for right forelimb markers during wake movements. Group average ± SEM is plotted in green; thin grey lines show individual animals (N = 7).
**(B)** Same as (A) for REM-sleep twitches, plotted in purple.
**(C)** Violin plot comparing peak displacement (movement amplitude) for all wake movements (green) and twitches (purple) pooled across animals (wake n = 4,365; twitch n = 3,766). The black line is the mean and the red line is the median.
**(D)** Polar scatter plot of every wake movement: angle denotes movement direction and radial distance indicates amplitude. The grey filled areas are a histogram illustrating the proportion of movements in each directional bin.
**(E)** Polar scatter plot of every twitch, formatted as in (D) but shown in purple, highlighting their smaller amplitudes and more evenly-distributed direction.
